# Claspin-dependent and -independent Chk1 activation by a panel of biological stresses

**DOI:** 10.1101/2022.11.29.518268

**Authors:** Hao-Wen Hsiao, Chi-Chun Yang, Hisao Masai

## Abstract

Replication stress has been suggested to be an ultimate trigger of carcinogenesis. Oncogenic signal, such as overexpression of CyclinE, has been shown to induce replication stress. Here, we show that various biological stresses, including heat, oxidative stress, osmotic stress, LPS, hypoxia, and arsenate induce activation of Chk1, a key effector kinase for replication checkpoint. Some of these stresses indeed reduce the fork rate, inhibiting DNA replication. Analyses of Chk1 activation in the cell population with western analyses showed that Chk1 activation by these stresses is largely dependent on Claspin. On the other hand, single cell analyses with Fucci cells indicated that while Chk1 activation during S phase is dependent on Claspin, that in G1 is mostly independent of Claspin. We propose that various biological stresses activate Chk1 either directly by stalling DNA replication fork or by some other mechanism that does not involve replication inhibition. The former pathway predominantly occurs in S phase and depends on Claspin, while the latter pathway, which may occur throughout the cell cycle, is largely independent of Claspin.

Our findings provide evidence for novel links between replication stress checkpoint and other biological stresses and points to the presence of unknown mechanisms of Chk1 activation in mammalian cells.

## Introduction

Genome instability is a major driving force for cancer development [1]. Oncogenic stress has been shown to induce replication stress, which is the trigger for induction of genome instability. How oncogenic stress (e.g., Cyclin E overproduction) causes replication stress is still not clear, but the reduced levels of cellular nucleotide pool induced by oncogenic stress were shown to cause genome instability [1–2].

Living organisms are exposed to various types of stress, and are equipped with a variety of systems to deal with them [2]. For example, in the cellular response pathway to replication failure, the stress signal is transmitted from sensor kinase (ATR) to effector kinase (Chk1) to temporarily arrest progression of replication and cell division. Claspin is involved in the ATR-Chk1 signaling axis in the replication stress response as an essential mediator [3–7].

Claspin and its yeast homologue, Mrc1, are essential for activation of downstream effector kinases (Chk1 and Cds1/Rad53, respectively) as replication checkpoint mediators [8–13]. Chk1 binding domain (CKBD) in metazoan Claspin were reported to be required for regulated Chk1 interaction [7]. It was also reported that Claspin could promote Chk1 activation in the presence of ATR in vitro [10]. Recently, we and others reported that either Cdc7 or CK1γ1 can phosphorylate CKBD of Claspin for checkpoint activation though to different extents depending on cell types [3, 14].

Cellular responses to environmental signals are important for cell proliferation and survival. Although detailed studies have been conducted on cellular responses induced by various types of stress, how these cellular responses cross talk and control cell proliferation and survival in an integrated manner has been largely unknown. Recently, it has been reported that DNA damages and/or Chk1 phosphorylation are induced by biological stresses, including ultraviolet (UV), arsenate (Ar), NaCl, lipopolysaccharides (LPS), hypoxia, heat shock, H_2_O_2_, and high glucose (HG) [15–24], suggesting the presence of cross talks between various biological stress responses and replication checkpoint.

Ar and arsenite are derivatives from arsenic. However, due to the stronger cytotoxicity of arsenite than that of Ar, arsenite has been more extensively studied than Ar. Arsenite has been shown to interfere with DNA repair machinery and induce apoptotic cell death through regulating ATR, Chk1, and Chk2 signaling pathway [18, 25–26]. Furthermore, 400 uM Ar treatment for 1 hr has also been demonstrated to activate integrated stress responses through important eIF2α kinases in MEFs [27].

Hypoxia was also reported to induce DNA damages and replication checkpoint activation through inducing expression of ATRIP, an activator of ATR [17, 28–29]. It was also observed that hypoxia activated unfolded protein response (UPR) which was sensed by PERK, IRE1 and ATF4 and that hypoxia partially blocked ongoing replication forks through PERK and decreased the capacity of new origin firing, suggesting that replication stress was generated by hypoxia [30–34]. Notably, Claspin-Chk1 axis negatively regulates DNA replication during UPR. All the malfunctioned replication phenotypes triggered by hypoxia contribute to replication catastrophe potentially through inducing the expression of APOBEC3B, a DNA cytosine deaminase, further disrupting genome stability [35].

Furthermore, LPS treatment (1 ng/ml, 1 hr) has been demonstrated to down-regulate the gene expression associated with mitosis, DNA replication, DNA repair and G1/S transition (e.g. Mcm2-5 and RAD51), in human and murine macrophages, and hypercapnia (high CO_2_ concentration; 20% CO_2_) was able to reverse this process [36]. Moreover, LPS treatment in combination with IL-4 induced Chk1 phosphorylation as well as DNA damage responses in B cells, although this may be due to the induction of CSR which involves double-stranded DNA breaks [35]. Therefore, bacterial LPS is a potential agent that affects DNA replication and induces replication checkpoint [36–37]. Moreover, heat-induced Chk1 activation, which depended on Rad9, Rad17, TopBP1 and Claspin, was reported in HeLa cells and chicken B lymphoma DT40 cells [38]. It has also been shown that ATR-Chk1 axis is preferentially activated in HCT116 cells and Jurkat cells, a human T cell leukemia cell line, in response to heat shock (42-45°C) and Chk1 inhibition in conjunction with heat shock can enhance apoptotic cell death [16, 39].

In response to osmotic shock (NaCl), budding yeast Mrc1, homologue of Claspin, was phosphorylated by Hog1 kinase, and early-firing origins were delayed [40]. This response, however, does not involve Mec1 (sensor kinase) or Rad53 (effector kinase). Consistent with the finding in yeast [40], Claspin is directly phosphorylated by p38 MAP kinase, mammalian homologue of Hog1 kinase, and safeguards cells from DNA damages elicited by osmotic stress [19–20].

On the other hand, oxidative stress/H_2_O_2_ produced reactive oxygen species (ROS) and posed replicative threats by inducing replisome disassembly, stalling replication forks and generating DNA breaks, comparable to the effect of HU [41–42]. Elevated ROS levels in response to H_2_O_2_ dissociated peroxiredoxin 2 (PRDX2) and Timeless from the chromatin, whose binding is critical for replication fork progression [42]. Moreover, the involvement of APE2, Apurinic/apyrimidinic (AP) endonuclease, which contains Chk1-binding motifs, is required for oxidative stress-induced Chk1 activation in a manner dependent on ATR in *Xenopus* egg extracts [24].

Additionally, HG condition can cause replication stress through provoking nucleotide imbalance. It introduces chemical modifications on DNA as a result of the adduct of anomalous glucose metabolism, giving rise to genome instability [22-23, 43-44]. Interestingly, HG (37.8mM glucose) compromised Chk1 activation and DNA damage response 1 hr after UV irradiation or etoposide treatments, suggesting HG condition confers radio- and chemoresistance in cells. However, whether HG conditions alone activate Chk1 is not clear [22].

These results strongly suggest that replication checkpoint can be activated upon a wide spectrum of cellular stresses to maintain genome integrity. However, how cellular responses to various biological stresses are linked to activation of replication checkpoint is largely unexplored [4].

We have reported novel functions and mechanisms of Claspin actions in replication initiation and in replication checkpoint activation [3-4, 45-46]. More recently, we reported that Claspin regulates growth restart from serum starvation by activating the PI3K-PDK1-mTOR pathway [45]. Here, we have examined a potential role of Claspin in replication checkpoint activation in response to various cellular stresses, and show that Claspin plays a crucial role in cellular responses to heat shock, hypoxia, arsenate, NaCl, oxidative stress, LPS and HG. We show that some stresses suppress DNA replication and others do not have much effect on it. The Chk1 activation occurs throughout cell cycle, but that outside the S phase is less dependent on Claspin than that within the S phase. We have concluded that various biological stresses activate Chk1 either by direct activation of replication checkpoint in a Claspin-dependent manner or through distinct pathways that is independent of Claspin. The results also points to the presence of unknown mechanisms of Chk1 activation in mammalian cells.

## Results

### Various biological stresses activate Chk1 and induce DNA damages

We examined the effect of various stresses on DNA damages and Chk1 activation by analyzing single cells through immunostaining (**Fig. 1A**). The biological stresses chosen were, in addition to HU and UV (replication stresses), high temperature (heat stress), NaCl (osmotic stress), Ar (arsenate salt), LPS (bacterial infection), H_2_O_2_ (oxidative stress), HG (high glucose) and hypoxia (hypoxic stress).

**Figure 1:**
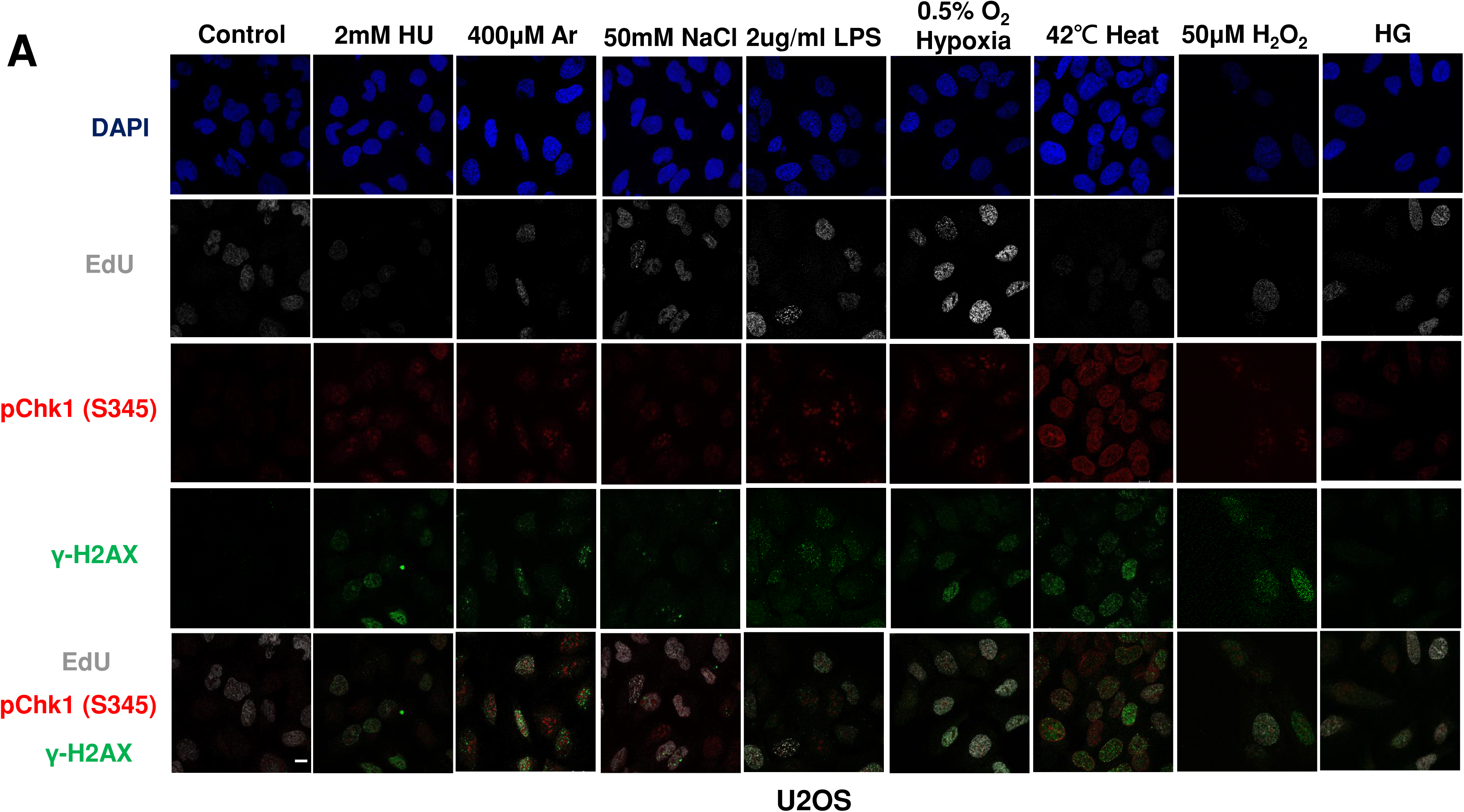
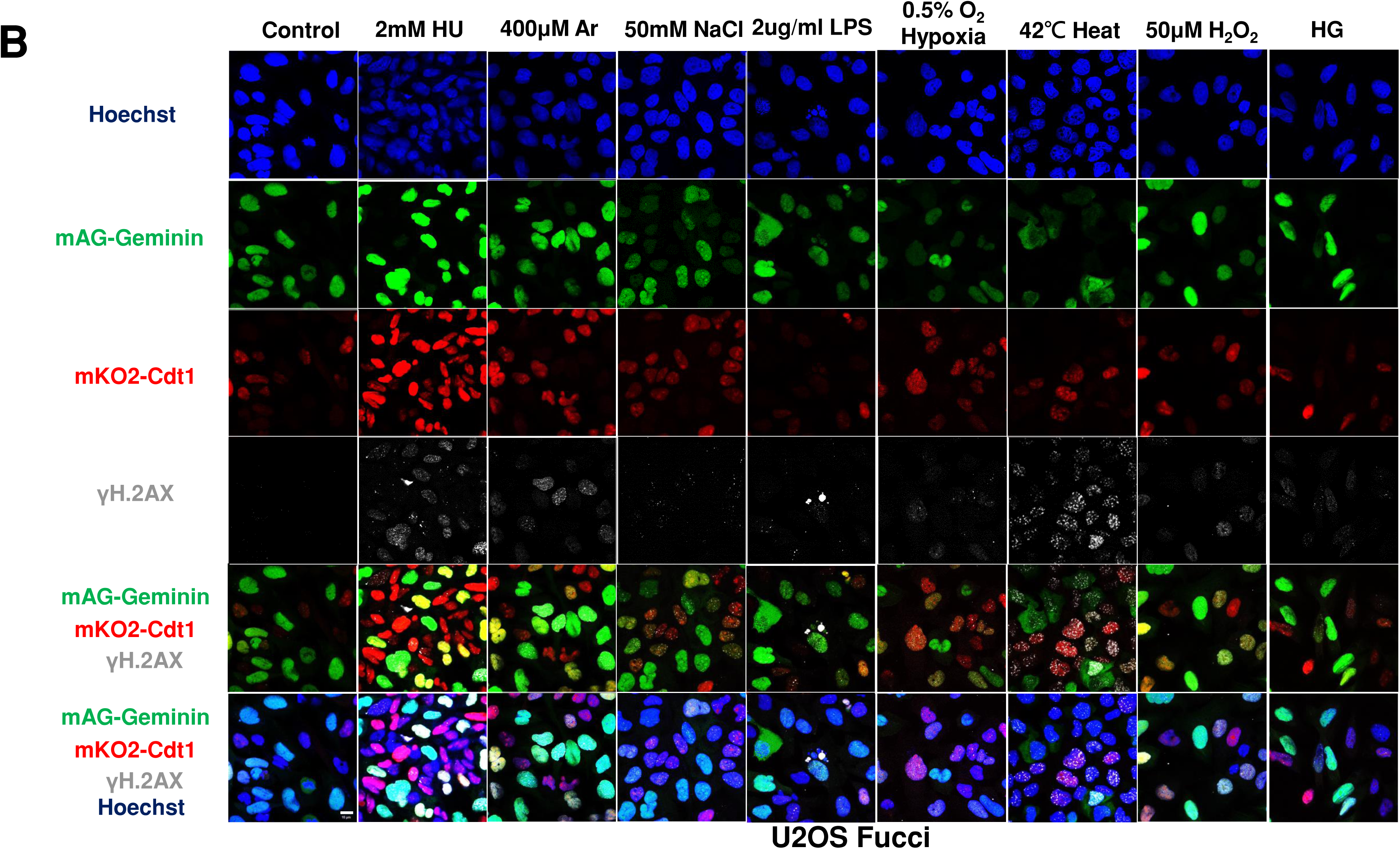
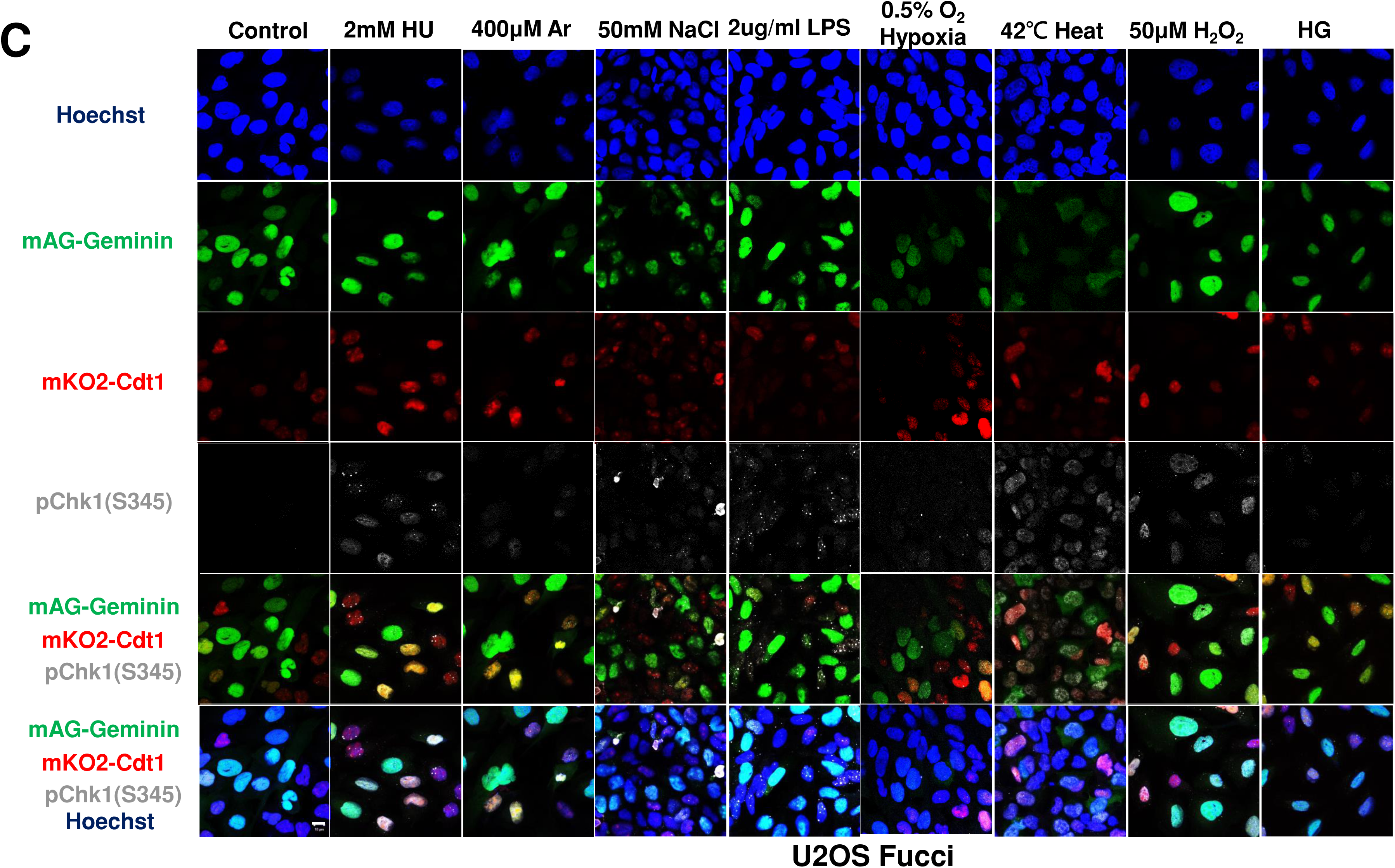
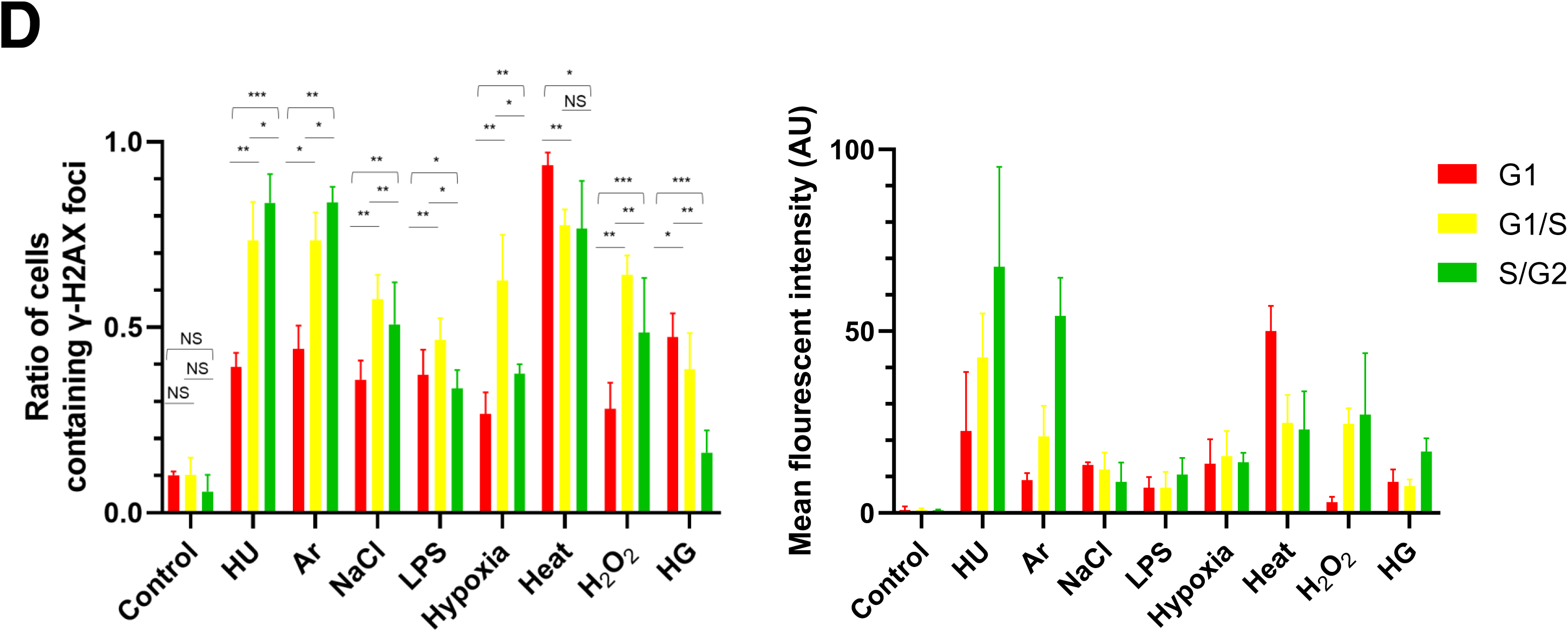
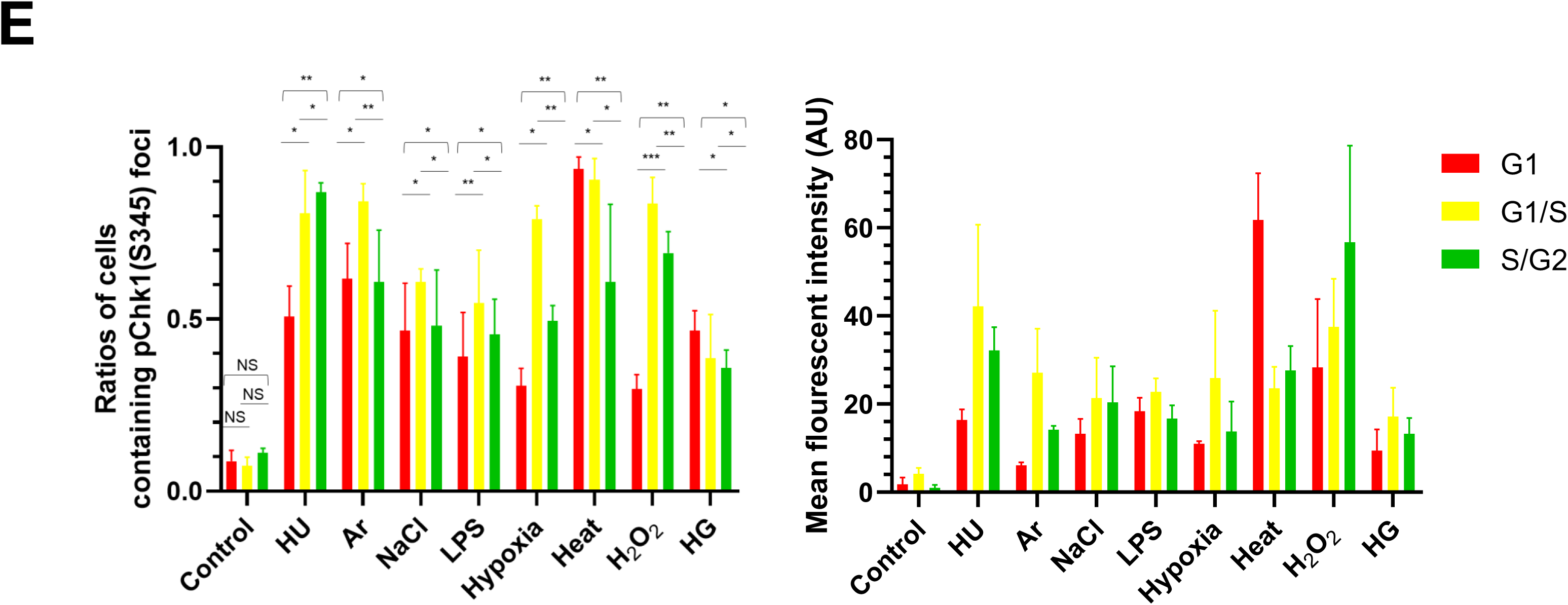
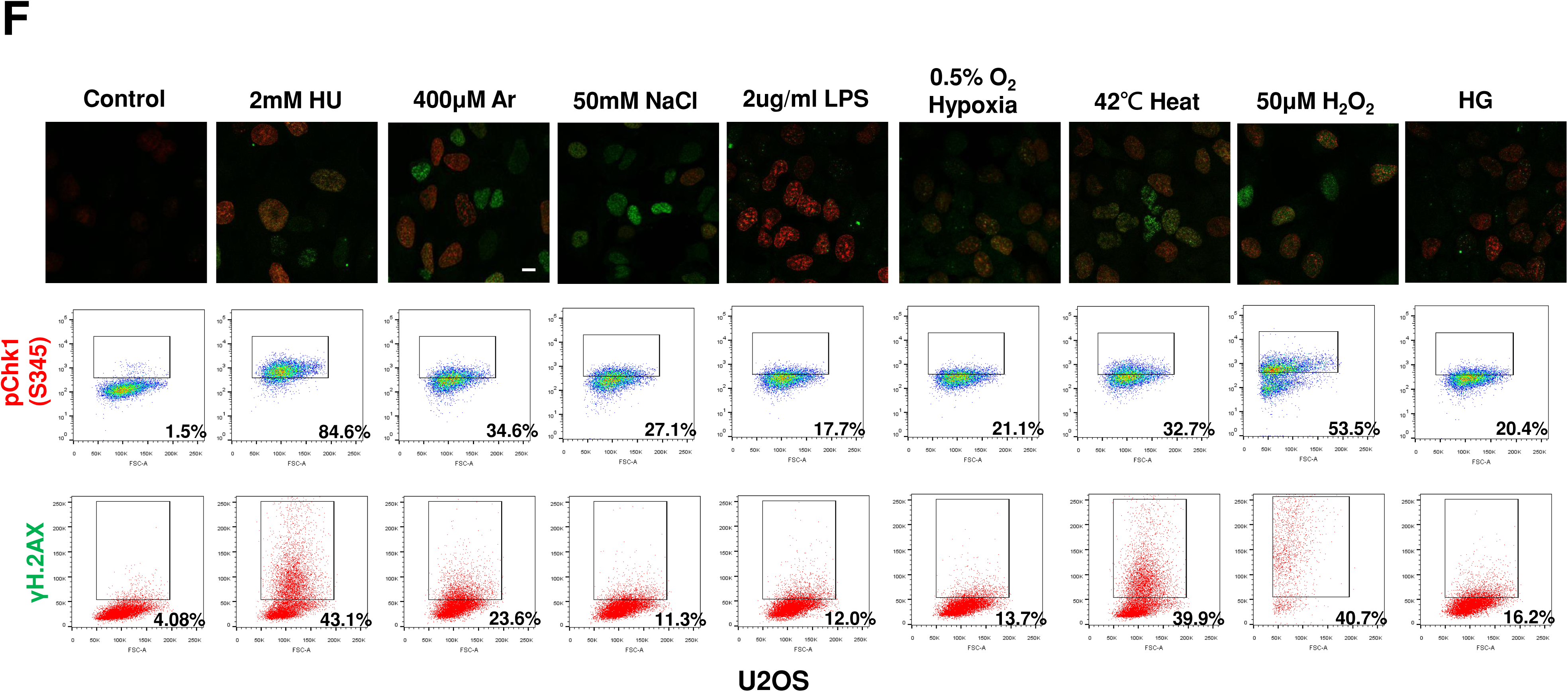
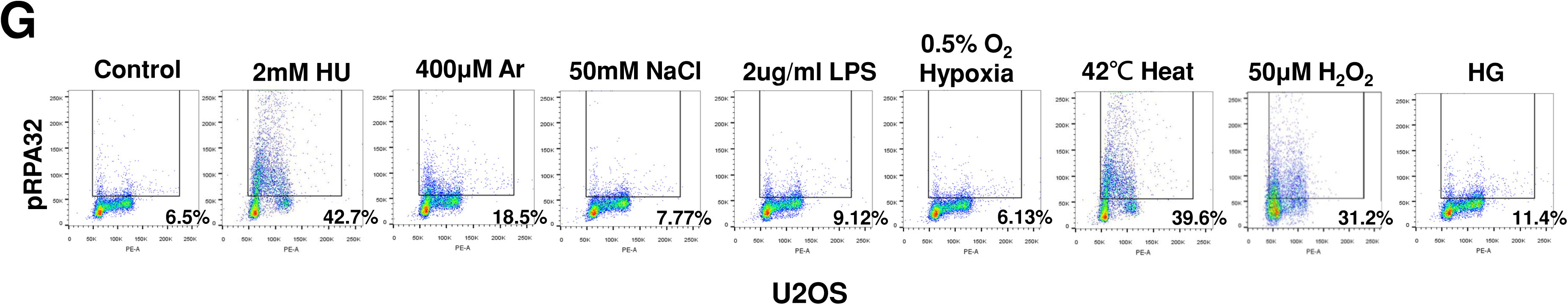
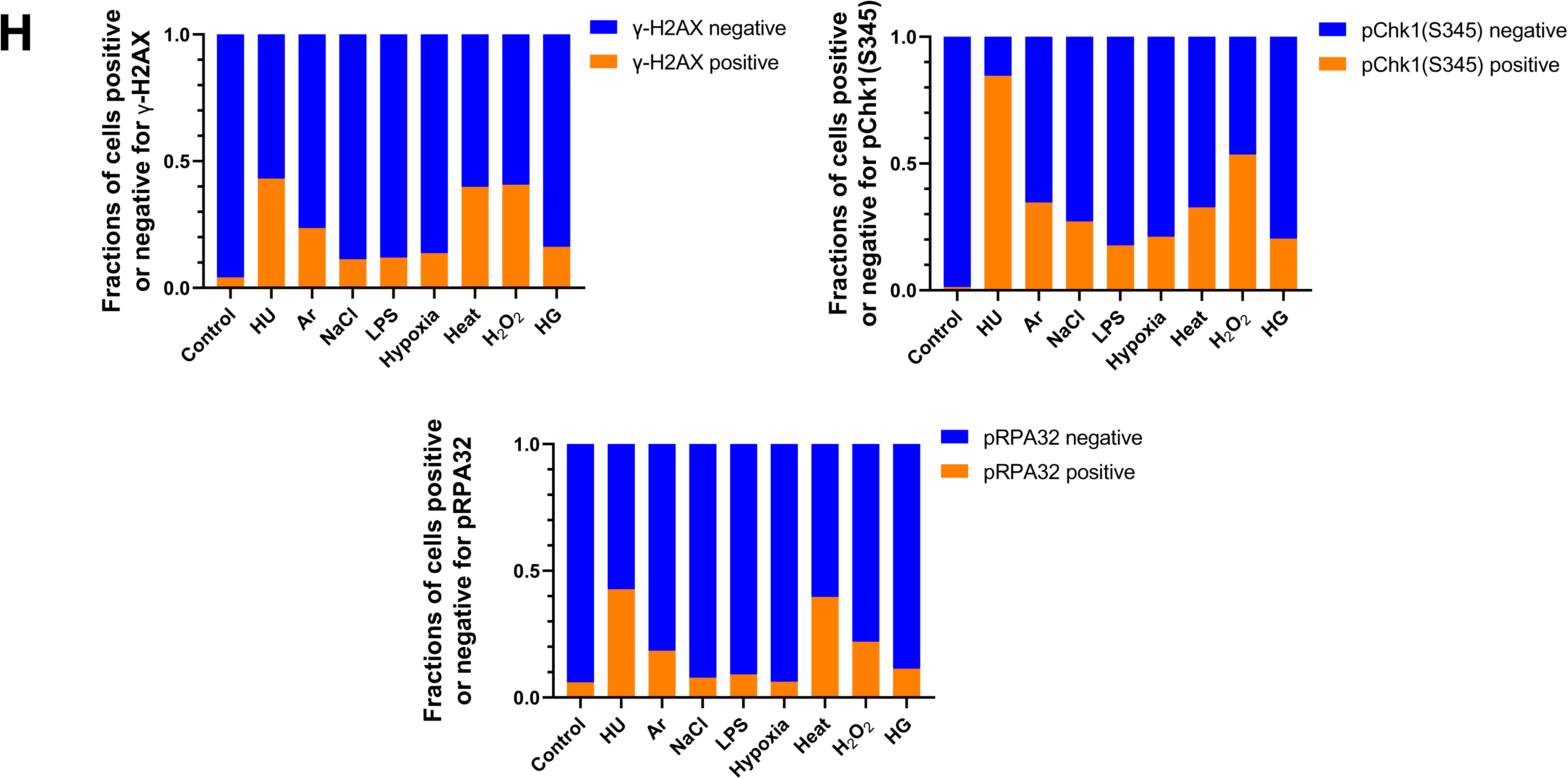
Various cellular stresses differentially affect DNA replication and induce DNA damages and replication checkpoint in a cell cycle stage-dependent manner. A. U2OS cells were exposed to indicated cellular stresses for 3 hr, EdU-labeled for 15 min and stained with indicated markers. Cells were then visualized and analyzed by confocal microscope Zeiss LSM780. Representative images are shown. Scale bar is 10 μm. Green, DAPI (DNA); white, EdU (DNA synthesis); red, pChk1(S345) (replication checkpoint); green, γ-H2AX(DSB). B and C. U2OS Fucci cells were exposed to indicated cellular stresses for 3 hr and subjected to immunostaining. Cells were then analyzed by confocal microscope Zeiss LSM780. Representative images are shown. Scale bar is 10 μm. Blue, Hoechst (DNA); green, geminin (S/G2 marker): red, Cdt1 (G1 marker); white, γ-H2AX; yellow in the merged image, G1/S boundary. D and E. Left: Fractions of U2OS Fucci cells containing γ-H2AX (D) and pChk1(S345) (E) foci were quantified for each cell cycle population. Right: The mean fluorescent intensity of γ-H2AX (D) and pChk1(S345) (E) was quantified for each cell cycle population. AU: arbitrary unit. F. U2OS cells were exposed to different stresses for 3 hr, and then were subjected to γ-H2AX (green) and pChk1(S345) (red) staining, followed by flow cytometry and confocal microscopy (Zeiss LSM780) analyses. Representative data and images are shown. Scale bar is 10 μm. G. Representative FACS data of pRPA32 (S4/8) staining (single-stranded DNA) of U2OS cells exposed to various stresses. H. Quantification of the data from F and G. Fractions of γ-H2AX, pChk1(S345) or pRPA32 (S4/8)-positive populations are indicated for cells exposed to various stresses. All statistical analyses represented the mean values ± SEM of indicated mean fluorescence intensity under two independent experiments, all of which included three replicates (∗p<0.05, ∗∗p<0.01, *** p<0.001, ns: no significant difference).

We noted that all the biological stresses used induced Chk1 phosphorylation at S345 (pChk1(S345)) to different extents after 3 hr treatment. Under the same condition, γ-H2AX foci appeared in most cells exposed to these stresses, albeit to different extents. We also noted that some stresses (Ar, heat, H_2_O_2_) greatly reduced EdU foci, suggesting their inhibitory effects on DNA replication (**Fig. 1A; see also** **Fig. 2**).

**Figure 2:**
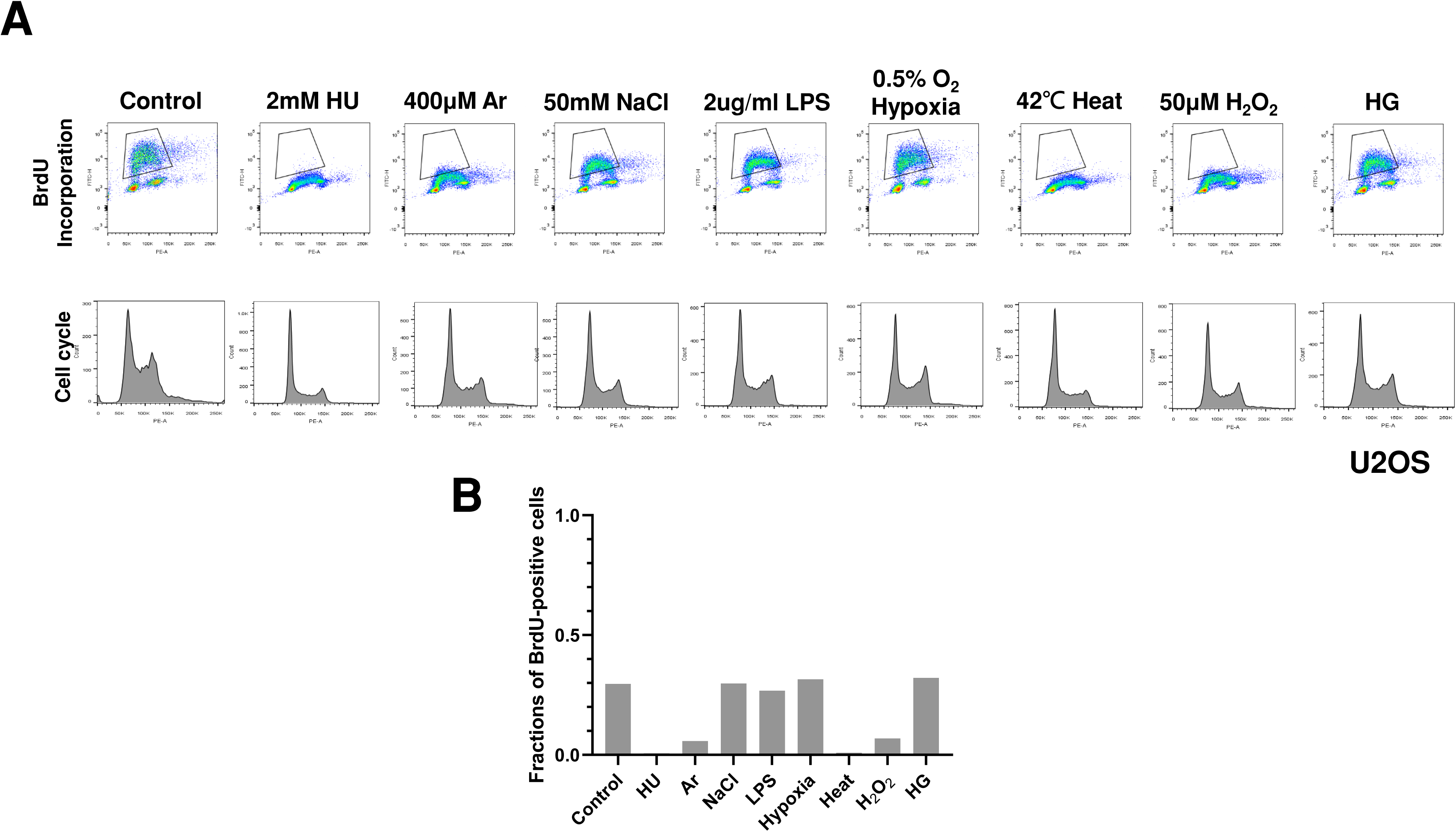
Various biological stresses could influence DNA replication rate to different extents and in a different time course manner. **A.** U2OS cells were treated with indicated biological stresses for 3hr. The nucleotide analog BrdU was added for 15 min before the cell harvest. Cells were then stained with anti-BrdU antibody and propidium iodide (PI) and analyzed by flow cytometry. Upper, BdU incorporation (DNA synthesis); lower, cell cycle (DNA content). **B**. Fractions of BrdU-positive cells in stress-treated cells (gated in A) were measured and presented.

Above results indicate that Chk1 activation and DNA damages appear to be induced in most cells by any of the stresses. To determine the cell cycle specificity of Chk1 activation and DNA damages more accurately, we next tried to quantify the fractions of γ-H2AX- and pChk1(S345)-positive cells in EdU-incorporating cells to access the S phase specificity of DNA damages and replication checkpoint activation induced by each stress. However, due to strong inhibition of DNA replication by some stresses, it turned out to be difficult to accurately determine the relationship between cell cycle and DNA damages/Chk1 activation. Therefore, U2OS Fucci (Fluorescent Ubiquitination-based Cell Cycle Indicator) cells were treated with indicated stresses (**Fig. 1B-E**). Fucci cells expressed two cell cycle marker proteins, mKO2-Cdt1 and mAG-Geminin, marking G1-phase cells in red, cells in G1/S boundaries in yellow, and S/G2-phase cells in green [47]. We discovered that most stresses induced pChk1(S345) and DNA damages (γ-H2AX) throughout the cell cycle; however, to different extents (**Fig. 1B-E**). Next, we quantified γ-H2AX-postive cells in each cell cycle stage. γ-H2AX-positive cells were defined as cells with more than 5 foci of γ-H2AX. HU, Ar, NaCl, LPS, Hypoxia and H_2_O_2_ induced γ-H2AX-foci more preferentially in cells in G1/S transition and in S phase; whereas heat and HG activated γ-H2AX also G1 phase to significant extents (**Fig. 1B-C**). On the other hand, HU, Hypoxia and H_2_O_2_ induced pChk1(S345)-foci preferentially in cells in G1/S boundaries and in S phase. Heat and HG induced pChk1(S345)-foci in G1 cells more efficiently than in S phase cells, whereas Ar, NaCl and LPS activated Chk1 in all the cell cycle phases to similar extent (**Fig. 1D-E**). Strikingly, heat triggered pChk1(S345) foci formation in approximately 95% of G1-phase cells. Heat also induces γ-H2AX foci in more than 90% of the G1 cells. Similarly, fractions of G1 cells that showed pChk1(S345) and γ-H2AX signals under HG conditions were also higher than those of G1/S boundary and S/G2 cells, although the fractions and intensities of the signals were lower than those of heat-induced ones (**Fig. 1D-E**). We have also calculated mean fluorescent intensity (MFI) of γ-H2AX and pChk1(S345) under stresses. The results revealed that HU, Ar and H_2_O_2_ induced stronger γ-H2AX MFI in G1/S boundary and S/G2 cells than in G1 cells, while NaCl, LPS, hypoxia, and HG exhibited similar levels of MFI of γ-H2AX throughout the cell cycle (**Fig. 1D**). Heat not only induced γ-H2AX foci in more than 90% of the G1 cells but also showed higher MFI of γ-H2AX in G1 cells than that in cells in G1/S boundaries and in G2 cells (**Fig. 1D**). Similarly, HU, Ar, and H_2_O_2_ showed more vigorous signals of pChk1(S345) preferentially in G1/S boundary, and S/G2 cells, while NaCl, LPS, and HG treatments showed similar levels of the signal intensity of pChk1(S345) throughout the cell cycle (**Fig. 1E**). Hypoxia exhibited stronger MFI preferentially in G1/S boundaries compared to G1 cells and S/G2 cells (**Fig. 1E**). Consistent with the result of cell numbers, heat exhibited higher MFI of pChk1(S345) in G1 cells, compared to G1/S boundary cells and S/G2 cells (**Fig. 1E**). Taken together, we show that different cellular stresses activated Chk1 phosphorylation and DNA damage signals to different extents, and the response was also differentially regulated during cell cycle. A notable conclusion is that all the stresses can activate Chk1 all through the cell cycle (**Fig. 1D and E**, left graphs). Generally, intensity of Chk1 activation in G1 cells is lower than that in G1/S/G2 cells (**Fig. 1D and E**, right graphs).

We next conducted FACS analyses to more accurately measure the DNA damages and Chk1 activation under different cellular stresses. Consistent with the results of immunostaining, HU, heat, and H_2_O_2_ (oxidative stress) induced γ-H2AX foci in nearly 40% of all the cells and arsenate salt (Ar) in 23.6% of the cells, while other stresses (NaCl, osmotic stress; LPS, bacterial infection; HG, high glucose; Hypoxia, hypoxic stress) induced γ-H2AX foci in approximately 11-16% (**Fig. 1F and H** and Table. 1). We also analyzed RPA32 phosphorylation at S4/S8 (pRPA32), a marker of DNA damage. 6.5% of the control cells without any treatment were pRPA32-positive, which could be due to spontaneous DNA damage during the ongoing DNA replication (**Fig. 1G and H**). HU, heat and H_2_O_2_ induced pRPA32-positive cells in, respectively, 42.7%, 39.6% and 31.2% population of all the cells, and Ar 18,5%; while other stresses induced pRPA32 only in 6.13-11.4% populations (**Fig. 1G and H** and Table. 1). We also analyzed pChk1(S345), and showed that HU, Ar, heat and H_2_O_2_ activated Chk1 in 84.6%, 34.6%, 32.7% and 53.5% of all the cells, while NaCl, LPS, Hypoxia, and HG induced pChk1(S345) in the 17.7∼27.1% population of all the cells (**Fig. 1F and H**). These results indicate that Ar, heat and H_2_O_2_ induce Chk1 activation and DNA damage signals, while other stresses induce DNA damage and pChk1 signals at a lower level, consistent with the results of single cell analyses.

**Table 1:**
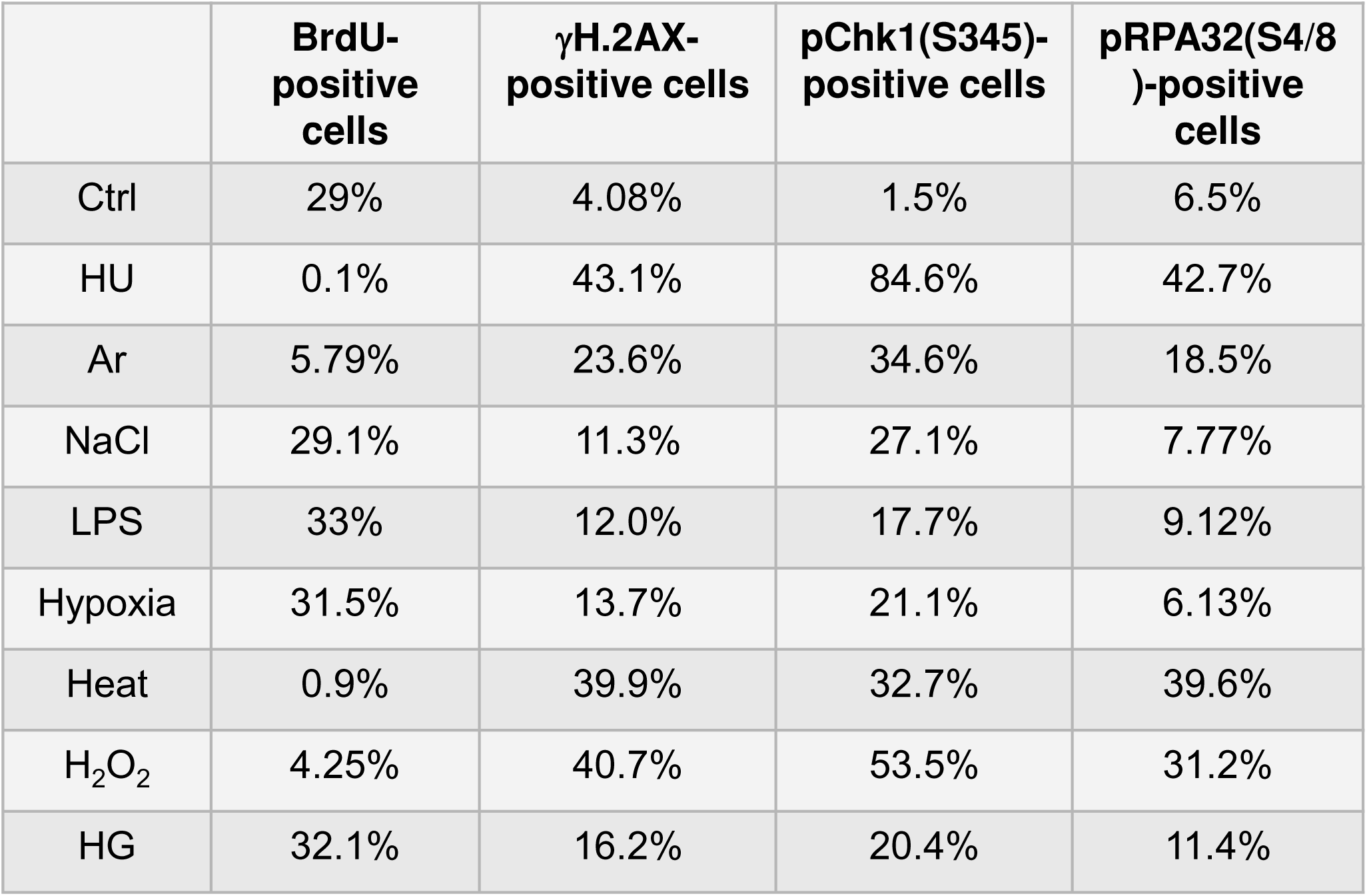
Fractions of cells positive for EdU, γ-H2AX, pChk1(S345), and pRPA32 in all the cell population in response to various biological stresses.

**Table 2:** Summary of pChk1(S345), γ-H2AX, pRPA32, and EdU in cells treated with various biological stresses. In the rows of “imaging”, the description is made on the basis of the data from the numbers of foci-positive cells (left panel of Figure 1D and E). “S>G1” indicates that foci are observed in S phase cells more frequently than in G1 phase.

### Biological stresses differentially affect DNA replication fork progression

To more precisely assess the effect of various biological stresses on DNA replication, we examined DNA synthesis in stress-treated cells by EdU incorporation assay (**Fig. 2A**). Consistent with the results of EdU imaging assay (**Fig. 1A**), HU, Ar, heat and H_2_O_2_ treatment for 3 hr greatly decreased BrdU incorporation, whereas other stresses did not significantly affect the BrdU incorporation (**Fig. 2A**). We also treated cells with various stresses for different periods of time and examined cell cycle and BrdU incorporation. Cell cycle profiles did not significantly change in HU, LPS, NaCl and heat treatment. On the other hand, H_2_O_2_ treatment for 24 hr led to increased G2 cell population, and UV treatment for 24 hr led to significant cell death **(data not shown)**.

We next conducted DNA fiber assays to examine DNA replication fork progression and determine replication fork speed under different stress conditions (**Fig. 3**). DNA was first labeled by CldU for 20 min, followed by IdU in the presence of various stresses for another 20 min (**Fig. 3A**). The ratio of IdU to CldU is the indicator of effect of the stresses on replication fork progression. The results showed that HU, heat, Ar and H_2_O_2_ significantly retarded replication fork progression, consistent with the reduced DNA synthesis shown by FACS and BrdU incorporation (**Fig. 2A**, **Fig. 3A**). On the other hand, other stresses did not significantly impede fork progression, consistent with the results of BrdU incorporation. These results indicate that, in addition to HU, Ar, heat and H_2_O_2_ inhibit DNA replication.

**Figure 3:**
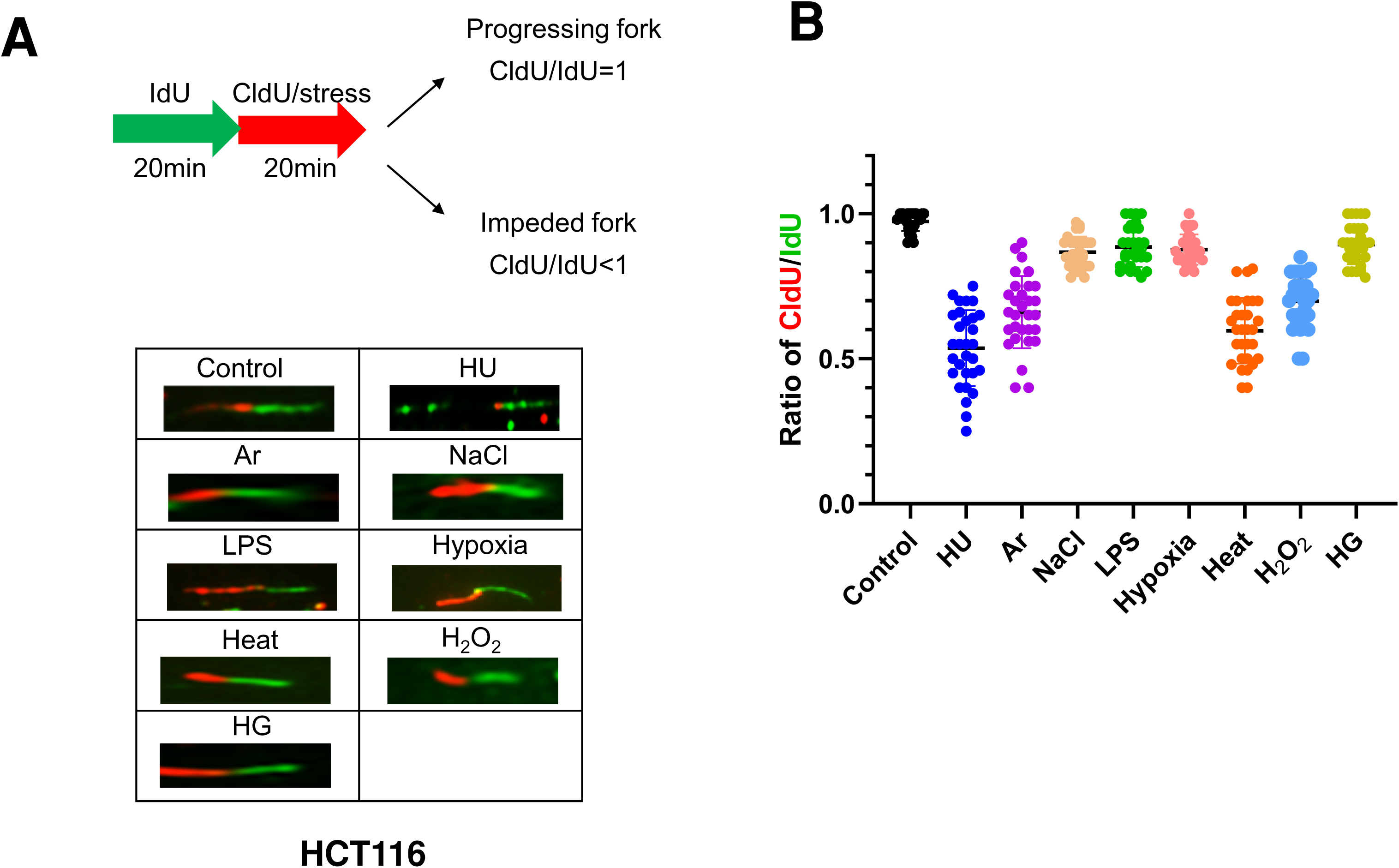
Different stress conditions differentially affect replication fork progression. **A**. Scheme for the experiments for monitoring replication fork progression. Briefly, HCT116 cells were labeled with IdU for 20 min, followed by the labeling with CldU in the presence of stresses for another 20 min. The ratios of CldU/IdU less indicate that replication fork progression is impeded by the stresses. Representative DNA fibers under different stress treatments are shown below the scheme. **B**. The CldU/IdU ratios in the presence of indicated stresses were determined and presented. All statistical analyses represented the indicated mean values ± SEM under two independent experiments.

### Effects of stresses on replication/ checkpoint factors

We then examined the expression of various factors by western blotting at 4 and 24 hr after different stress treatments. HU and UV strongly induced pChk1(S345) at 4 hr, while other stresses including heat, H_2_O_2_, NaCl, and LPS also induced pChk1(S345), albeit at a lower level. At 24 hr after the exposure to heat, pChk1(S345) was reduced to the non-stimulated level, suggesting that cells might already have recovered from the stress or have adjusted to the stress (**Fig. 4**). UV for 24 hr also led to loss of pChk1(S345) signal, but this was due to cell death induced by UV (see next section). In contrast, pChk1(S345) was still detected at 24 hr after treatment with HU, H_2_O_2_, NaCl and LPS.

**Figure. 4:**
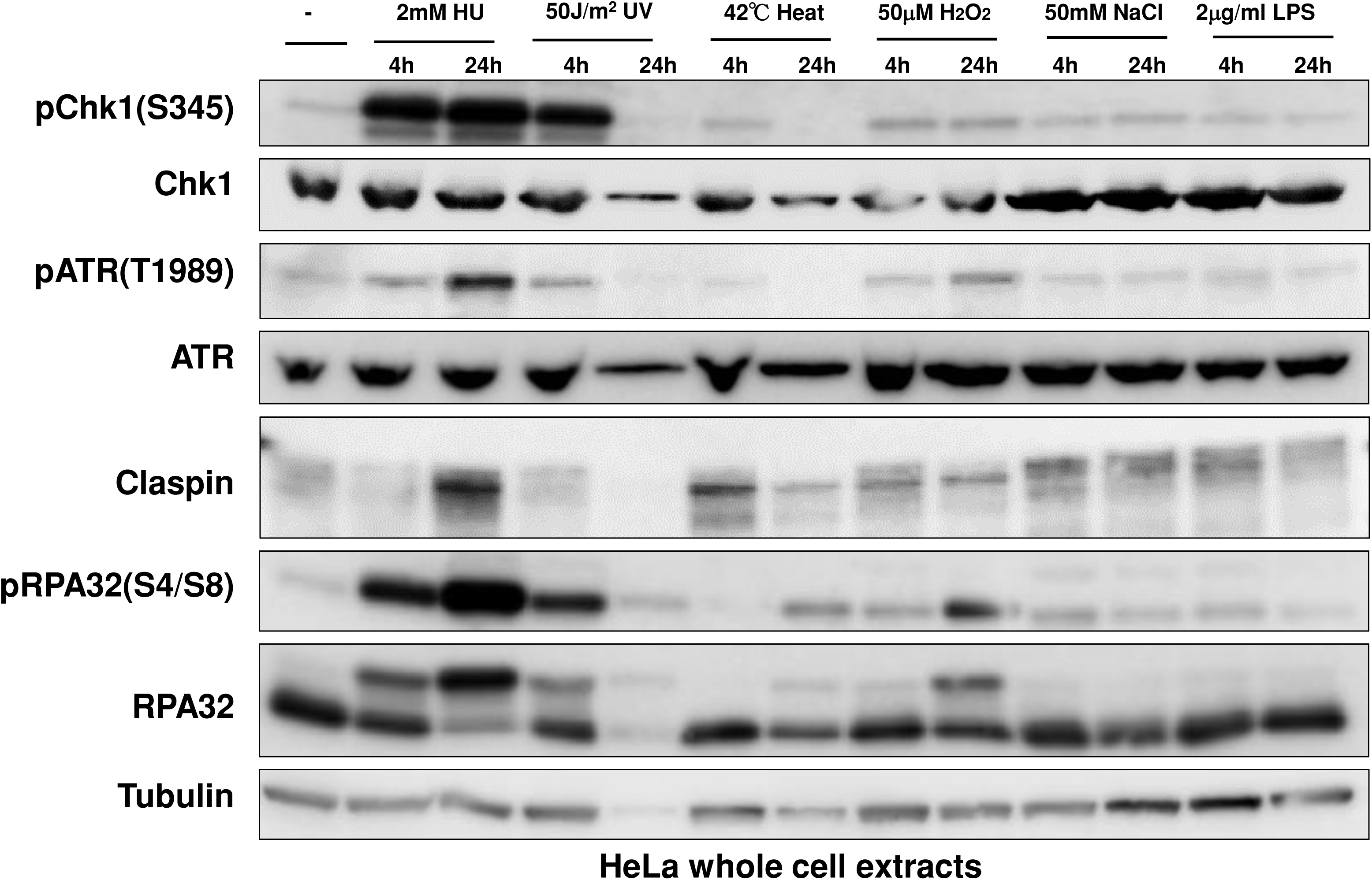
Effect of various biological stresses on checkpoint- and DNA damage-related factors in HeLa cells. HeLa cells were treated with indicated stresses for the time indicated. The whole cell extracts were analyzed by western blotting with the antibodies indicated.

ATR, the upstream PIKK (Phosphatidylinositol 3-kinase-related kinase), is required for Chk1 activation. Phosphorylation of ATR at T1989 is an indicator of ATR activation. ATR was activated not only by HU and UV, but also by H_2_O_2_, salt and LPS, albeit at a lower level. Heat slightly activated ATR at 4 hr but not 24 hr, similar to pChk1. Claspin undergoes phosphorylation upon replication stress (HU and UV), but also by other stresses, as exemplified by the mobility-shift on PAGE. It appears that Claspin undergoes differential phosphorylation upon various stresses, as suggested by differential mobility shift. RPA is phosphorylated at 24 hr by heat and H_2_O_2_, suggesting the induction of DNA damages by these stresses.

### Activation of bulk Chk1 kinase by various stresses depends on Claspin

Above results convincingly show that various biological stresses can activate Chk1 phosphorylation. Mobility-shifts of Claspin induced by these stresses suggest activation of Claspin during the processes. Using Claspin MEF^f/-^ cells that we previously established, Claspin can be knocked out by infection of *Ad-Cre* viruses. By using this cell line, we analyzed the requirement of Claspin for Chk1 activation by various stresses. Consistent with the results from HeLa cells, not only HU- or UV-treatment but also various stresses including heat, H_2_O_2_ and LPS, induced pChk1(S345) in MEF cells. In accordance with the requirement of Claspin for efficient phosphorylation of Mcm by Cdc7, Mcm2 phosphorylation was reduced by Claspin knockout (**Fig. 5A**). Chk1 phosphorylation, induced by various stresses, was not observed in Claspin knockout conditions (after *Ad-Cre* infection; **Fig. 5A**). Treatment with 50 mM NaCl induced only a low level of Chk1 phosphorylation (**Fig. 5A, lane 13**). 100 µM thymol (a phenol that is a natural monoterpene derivative of cymene and a volatile oil component) induced strong cell death, which was almost completely rescued by Claspin KO (**Fig. 5A, lanes 23 and 24**), indicating that thymol-induced cell death of MEF cells depended on Claspin.

**Figure 5:**
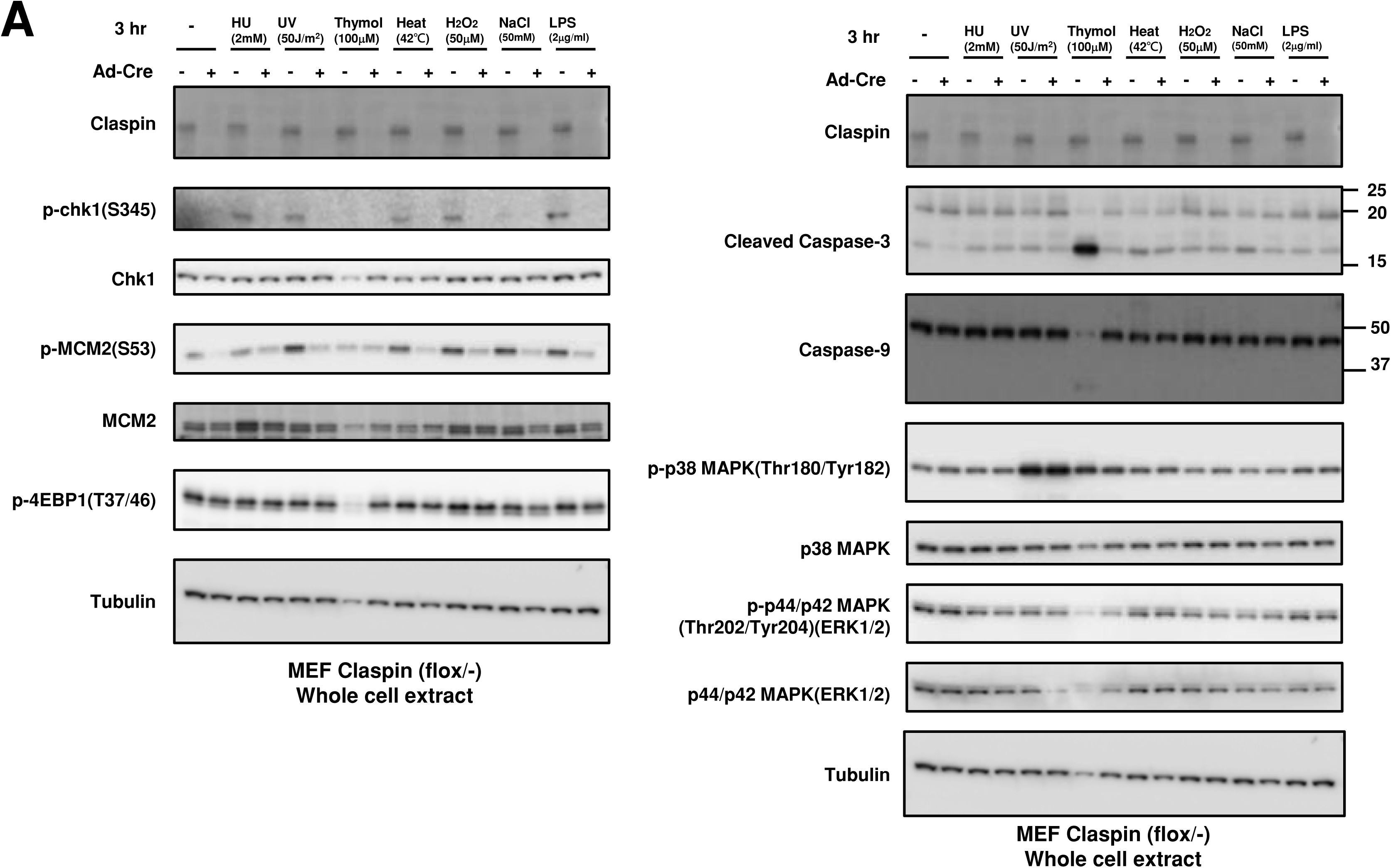
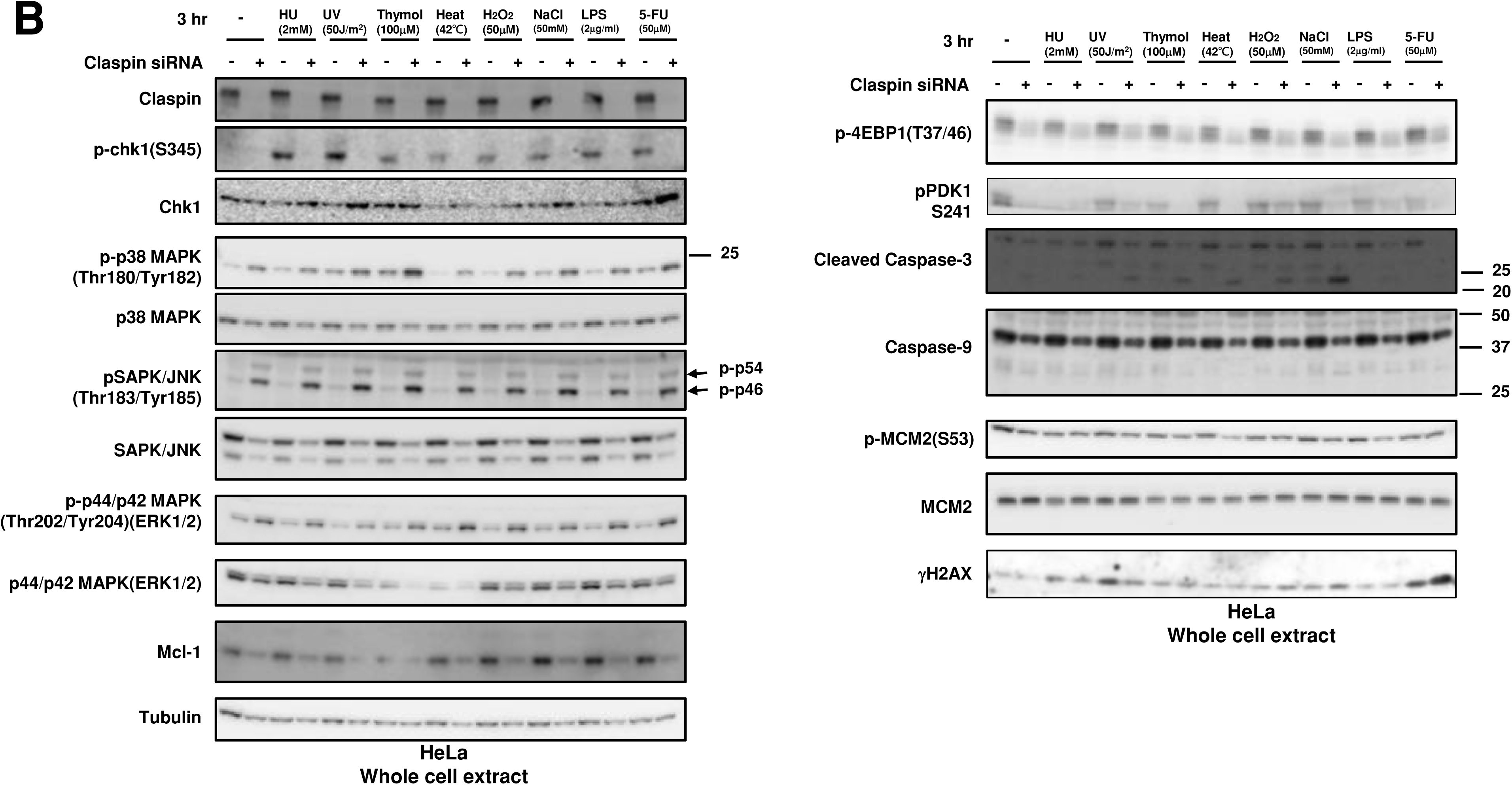
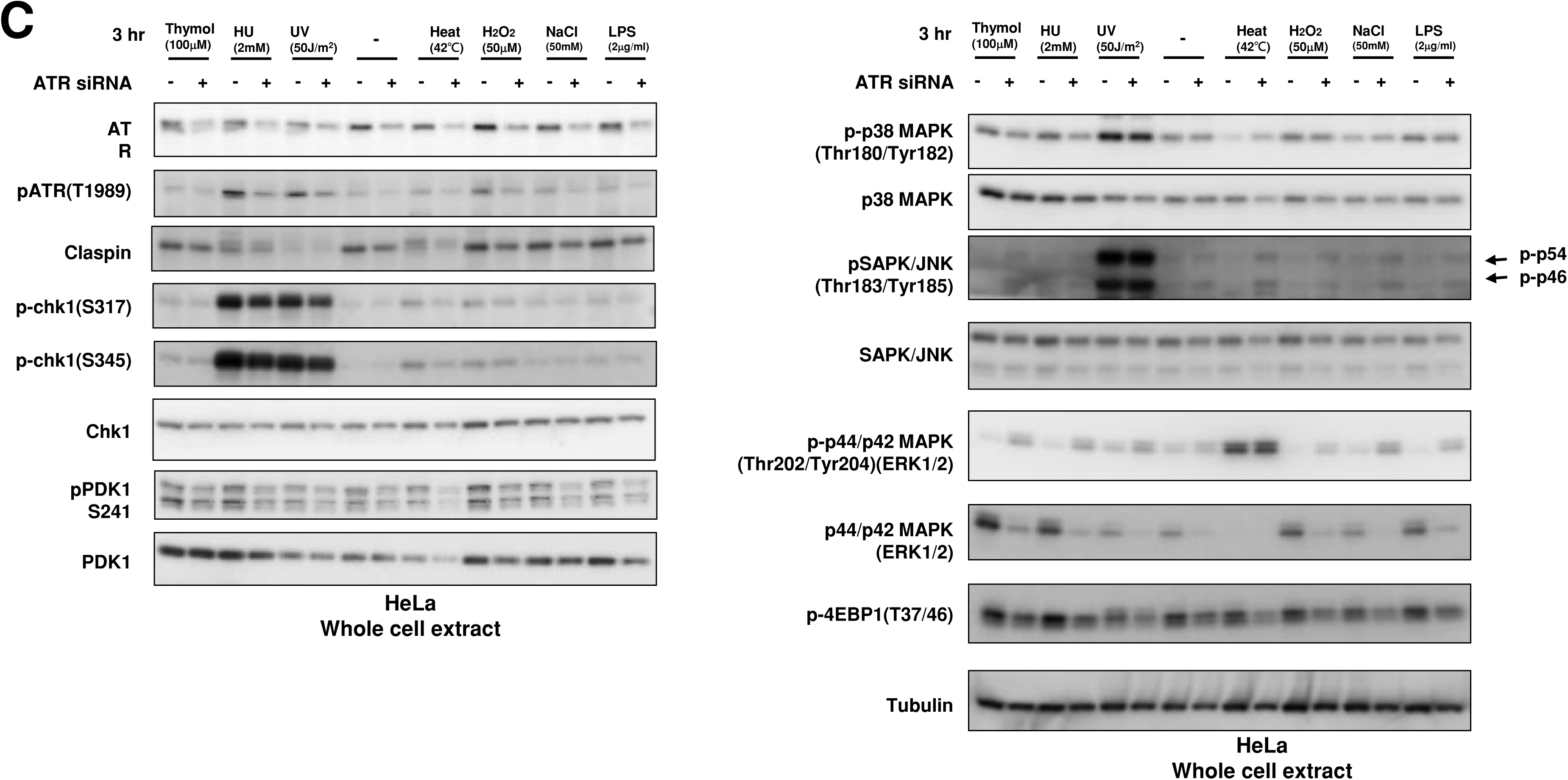
Effects of Claspin and ATR depletion on Chk1 activation by various biological stresses and on other factors involved in various growth-stimulated pathways. A, *Claspin(f/-)* MEF cells were treated with Ad-Cre or non-treated and exposed to various stresses for 3 hr. B. and C. HeLa cells were transfected with siRNA for Claspin (B) or ATR (C) for 24 hr (+) and were exposed to indicated stresses for 3 hr before the harvest. The whole cell extracts were analyzed by western blotting with antibodies indicated. -, control siRNA.

In HeLa cells, human cervical cancer cell line, the effects of Claspin siRNA on Chk1 activation by various stresses were examined. The same set of biological stresses activated Chk1 in a manner dependent on Claspin, although the levels of Chk1 activation were less than those achieved by HU or UV treatment (**Fig. 5B**). Notably, Claspin was mobility-shifted by all stresses examined in HeLa cells, as was observed in MEF cells (**Fig. 4**), but the extent and patterns of the shifts varied, suggesting the induction of different phosphorylation patterns of Claspin by different stresses (**Fig. 5B** and **C**).

We then examined the involvement of ATR, the upstream PIKK. ATR phosphorylation was induced by most of these stresses to differential extents, most notably by HU, UV and H_2_O_2_ (**Fig. 5C, lanes 3-6, 11,12**). ATR siRNA reduced Chk1 phosphorylation in cells treated with stresses (heat, H_2_O_2_ and LPS), suggesting ATR is required for Chk1 activation by some of the stresses (**Fig. 5C, lanes 9-12, 15 and 16**). The results also suggest a possibility that there could be other PIKKs that may be activated and transmit signals to Claspin. In summary, western analyses of Chk1 activation in the cell population indicate Claspin is required for Chk1 activation by these varieties of biological stresses, while ATR also plays a role for Chk1 activation at least by some of the stresses.

### Roles of Claspin in regulation of MAP kinase cascade and the PI3K-PDK1-Akt-mTORC1-4EBP1 pathway

In MEF cells, p38 MAPK or p44/p42 MAPK (ERK1/2), activated by MEK1/2 or MKK, respectively, was not affected by stresses or by Claspin depletion, except that UV treatment activated p38 MAPK (**Fig. 5B**, lane 21 and 22).

In HeLa cells, MAP kinases including p38 MAPK (Tyr180/Tyr182 phosphorylation), SAPK (stress-activated protein kinase)/JNK (Tyr183/Tyr185 phosphorylation) and p44/p42 ERK1/2 (Tyr202/Tyr204 phosphorylation) were activated by loss of Claspin (**Fig. 5C**, lanes 1-18), whereas the protein levels of these MAP kinases were generally slightly reduced by Claspin KD. The stresses did not alter the levels of these phosphorylated proteins with or without Claspin siRNA, except that UV and thymol slightly activated p38 MAPK (**Fig. 5C**, lanes 5 and 7).

4EBP1 (eukaryotic translation initiation factor 4E-binding protein 1) is known to be phosphorylated by mTORC1 in response to growth stimulation, and this phosphorylation is required for its release from eIF4E and subsequent activation of cap-dependent translation. PDK1 kinase is activated by PIP3, resulting in activation of Akt and the PKC isoenzymes p70 S6 kinase and RSK.

Our results illustrated that T37/46 phosphorylation of 4EBP1 was not affected by any stresses or by depletion of Claspin in MEF cells (**Fig. 5A**). In contrast, in HeLa cells, it was downregulated by Claspin knockdown, but not affected by any stresses examined. S241 phosphorylation of PDK1 was also inhibited by Claspin KD in HeLa cells, and was reduced by some stresses including HU, 5FU and thymol (**Fig. 5B**, lanes 21, 25, and 35). On the other hand, the Mcm2 phosphorylation was not affected under the same condition, as reported before. Similar effects were observed in other cancer cell lines, including U2OS and 293T cells **(data not shown)**. Weak cell death was induced by some stresses including UV, thymol, heat. H_2_O_2_, and salt in the absence of Claspin in HeLa cells, as indicated by the cleavage of Caspase-3 (**Fig. 5B**, lanes 24,26,28,30, and 32). The level of Mcl1, a member of Bcl2 protein family associated with anti-apoptotic activity, was reduced by Claspin KD (**Fig. 5B**).

Taken together, the results indicated that, in HeLa cells growing in the absence of stresses, Claspin plays suppressive roles in activation of the MAP kinase pathways, while it is required for activation of the PI3K-PKD-mTOR pathway

### Claspin-dependent and -independent activation of Chk1 by varieties of biological stresses

pChk1(S345) was induced by varieties of stresses not only in S phase cells but also in G1 phase cells (**Fig 1C and 1E**). We wondered if Claspin is required for Chk1 phosphorylation all through the cell cycle. To examine this, we used U2OS-Fucci cells and knocked down the expression of Claspin by siRNA, which was validated by western blotting (**Fig. 6A**). We quantified the fractions of cells showing pChk1(S345) signals under indicated cellular stresses in different cell cycle stages (**Fig. 6B and 6C**). We discovered that Claspin knockdown attenuated pChk1(S345) in S/G2 cells by 55 to over 70%, but it decreased pChk1(S345) in G1 phase cells only by 4 to 30 % under all the stress conditions except for NaCl (**Fig. 6D**). With salt stress, Chk1 activation was downregulated by ∼40% in both G1 and S phase cells. The results indicate that various biological stresses activate Chk1 all through the cell cycle but Claspin is required for Chk1 activation more predominantly during S phase.

**Figure 6:**
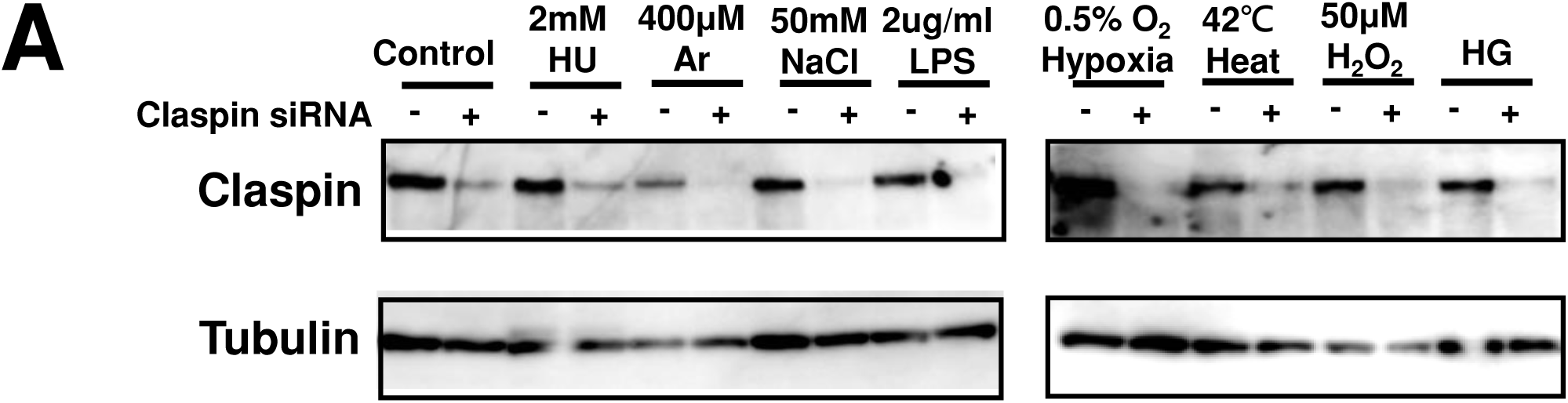
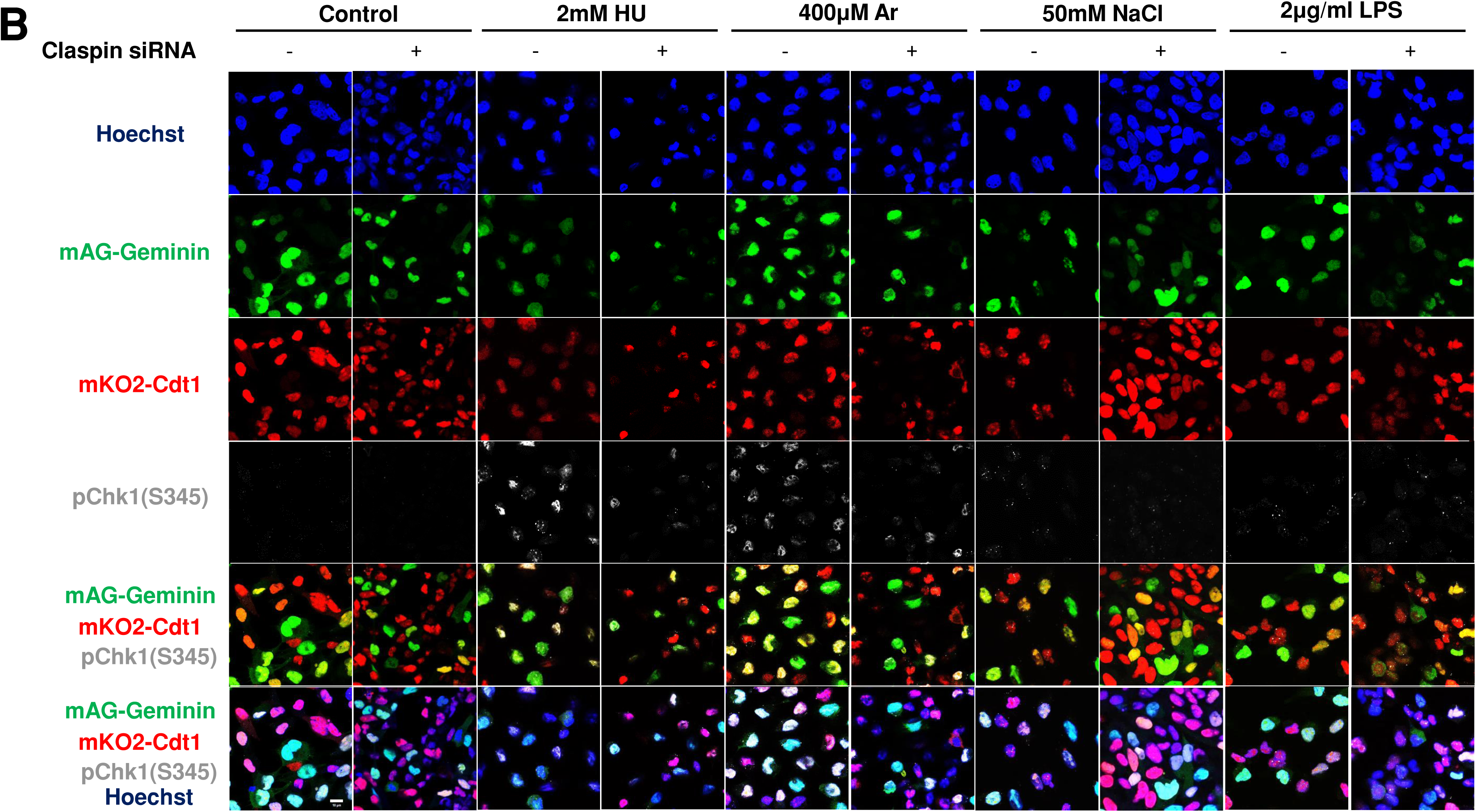
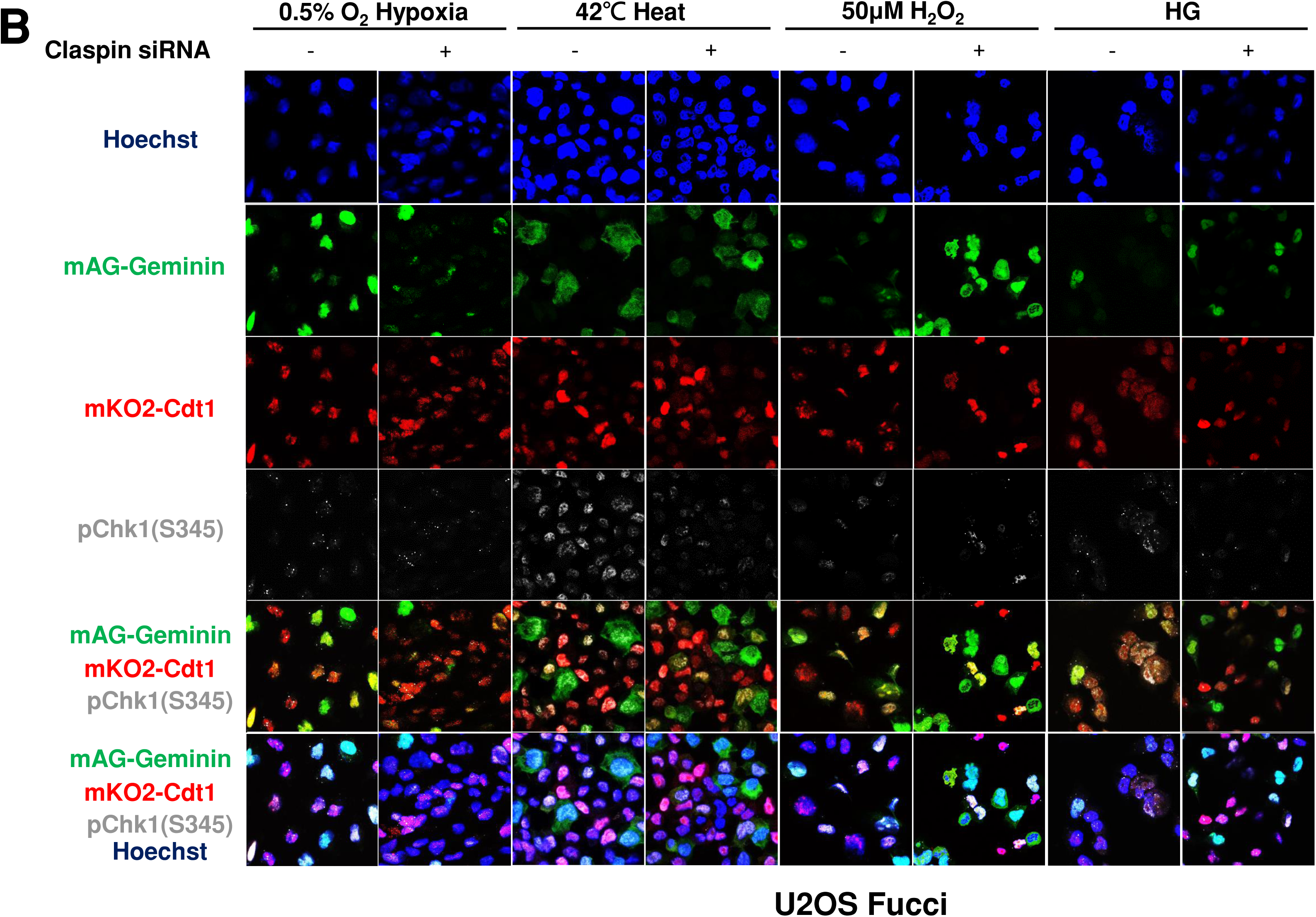
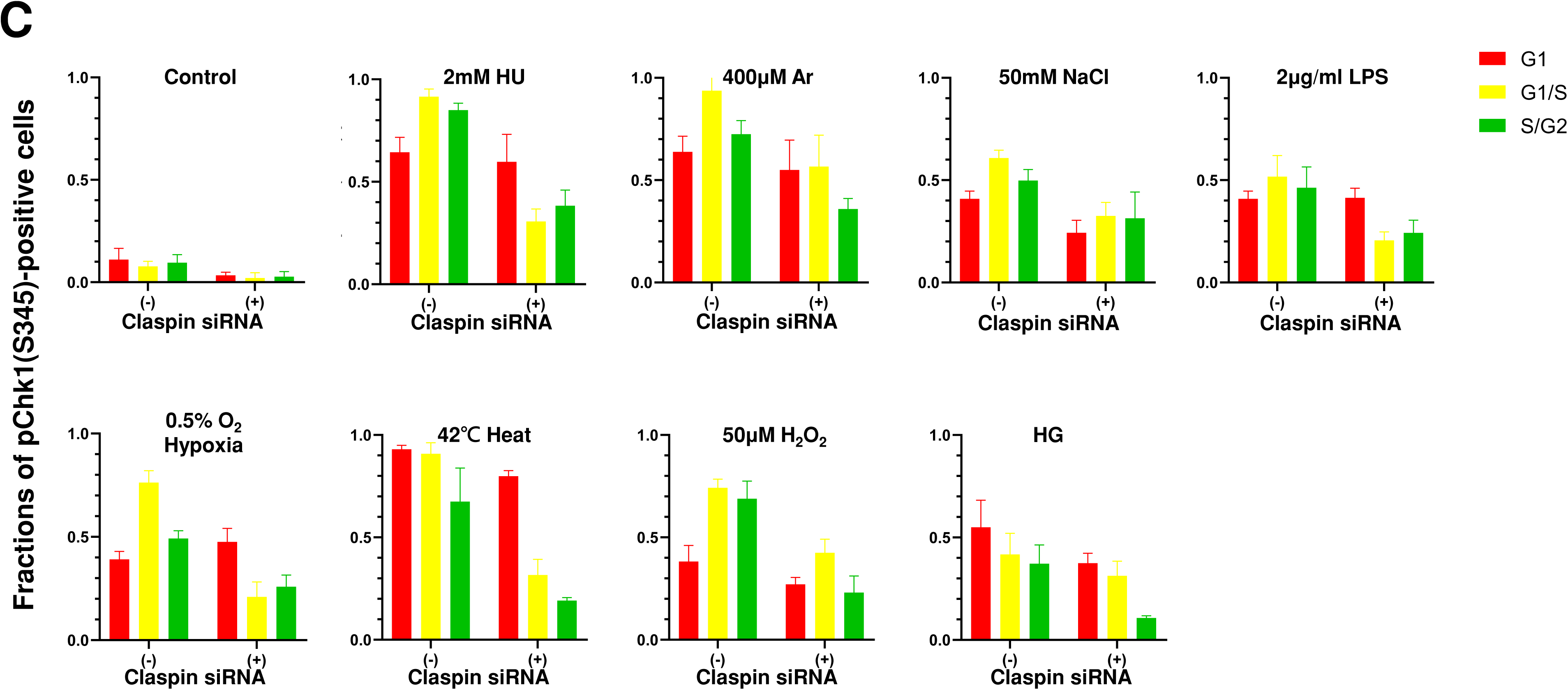
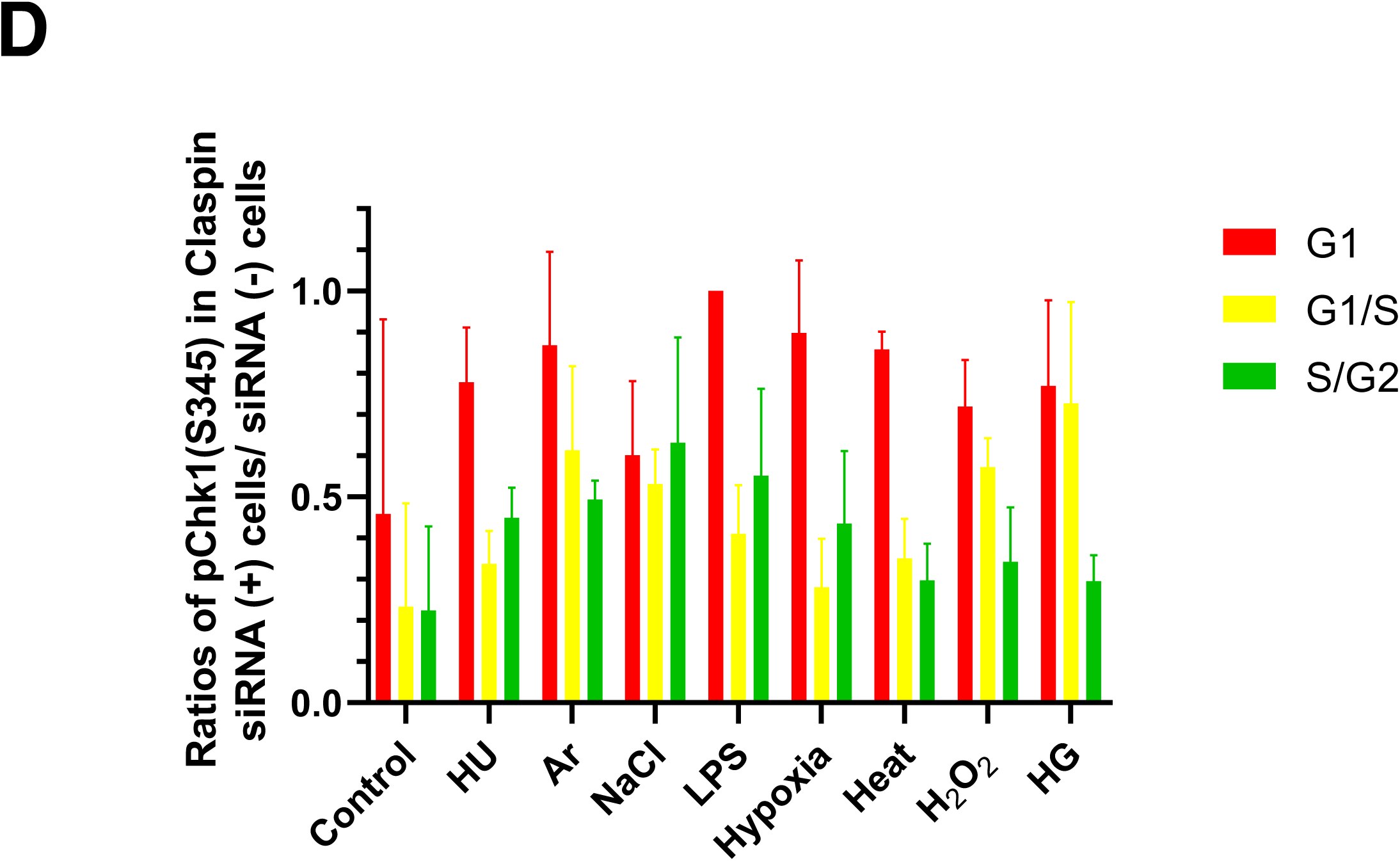
Claspin depletion abrogates Chk1 activation induced by various stresses mainly during the S phase. **A and B.** U2OS Fucci cells were transfected with Claspin siRNA or with control siRNA for 48 hr and were exposed to various stresses for 3 hr before the cell harvest. The whole cell extracts from a portion of the cells were analyzed by western blotting to detect Claspin and tubulin. **B**. The same cells were observed under confocal microscope Zeiss LSM710. Blue, Hoechst (DNA); green, mAG-Geminin (S/G2 cells); red, mKO2-Cdt1 (G1 cells); white, pChk1(S345) (replication checkpoint). **C.** Fractions of pChk1(S345)-positive cells in the U2OS Fucci cells of a specific cell cycle stage after exposure to various biological stresses. The signals were quantified by image J software. (+) and (-) refer to the cells transfected with Claspin siRNA and those transfected with control siRNA, respectively. **D.** Ratios of pChk1(S345)-positive cells in Claspin-depleted versus control cells in cells of the specific cell cycle stage. The smaller values indicate the less dependency of the pChk1 signal on the Claspin function. All statistical analyses represented the indicated mean values ± SEM under two independent experiments, all of which included three replicates (∗p<0.05, ∗∗p<0.01, *** p<0.001, ns: no significant difference).

## Discussion

Cells are equipped with various stress response pathways that protect cells and living species from various environmental stresses. Among them, replication stress is mostly observed during S phase by varieties of treatment that impede progression of replication forks. Previous studies have indicated that “oncogenic stress” triggers cancer cell formation through inducing replication stress. Replication fork stalling caused by varieties of oncogenic stress generates DNA damages, which eventually lead to accumulation of genetic lesions, causing tumors to be formed. Although experimental “oncogenic stress” includes overexpression of Cyclin E, E2F or growth factor receptors that can cause untimely growth stimulation, the nature of intrinsic “oncogenic stress” is rather unclearly defined.

### Biological stresses, DNA replication, Chk1 activation, ATR-Claspin and other signaling pathways

We here provide evidence that diverse stresses, including oxidative stress (H_2_O_2_), heat shock, osmotic stress (high salt), and LPS as well as arsenate, high glucose and hypoxia can activate Chk1. It appears that these stresses could be classified in two categories (**Supplementary Table S1**); one that arrests the replication fork and the other that does not obviously affect replication progression. The former may directly activate replication checkpoint, while latter may indirectly activate it. We show that in both cases, Claspin is required for Chk1 activation. We also showed that ATR may be required for Chk1 phosphorylation by these signals, although we cannot rule out the possibility that other PIKKs play a role.

It should be noted that there may be some discrepancies between our results and other previous published studies. For example, our finding that hypoxia did not drastically impede replication fork progression was somewhat contradictory to the one which showed hypoxia significantly retarded S-phase progression [17, 28–33]. Previous reports suggest inhibition of DNA replication by hypoxia treatment in RKO cells (poorly differentiated colon carcinoma cell line), but our DNA fiber and FACS analyses in U2OS or HCT116 cells showed no significant effect on replication fork progression or DNA synthesis (**Fig. 2 and 3**). This could be due to differences in the hypoxia condition. The concentration of Oxygen was 0.5% for 20 min for DNA fiber and 3 hr for FACS analyses in our experiments, in contrast to 0.1%, 8 hr in the previous report. Inhibition of DNA replication by hypoxia may require duration of low oxygen state for more than 3 hr.

In our assays, some stresses (HU, Ar, heat, and H_2_O_2_) can efficiently arrest replication forks; whereas other stresses (NaCl, LPS, hypoxia, and HG) do not (**Fig. 2 and 3**), and generally, those stresses that inhibit DNA replication also induce DNA damage signals (γ-H2AX and pRPA32). A previous study in HeLa cells showed that heat treatment for 2 hr in HeLa cells did not exhibit significant RPA32 phosphorylation [38]. Our western analyses show also that RPA32 phosphorylation is detected at 24 hr but not at 4 hr after heat treatment (**Fig. 4**). Thus, effects of various stresses on DNA replication and DNA damages could be affected by their strength and duration, as well as cell type used for the studies.

ATR activates two pathways; one leads to activation of Chk1 and the other to p38 MAP kinase [48]. Claspin is required for the former pathway, but not the latter. Claspin knockdown increased phosphorylation of MAP kinases including p38 MAPK, SAP1/JNK1, ERK1/2 in cancer cells, suggesting it may negatively regulate the MAP kinase pathways during unperturbed growth. We also showed that Claspin is potentially required for activation of the PI3K-PDK1-mTOR pathway. We recently demonstrated that Claspin is required for growth restart of serum-starved cells, and this is due to its essential role for activation of the PI3K-PDK1-mTOR pathway [45]. Thus, Claspin may play a role for the activation of this essential signaling pathway during normal growth of cancer cells.

### Chk1 activation during S phase depends on Claspin, but that in G1 is less dependent on Claspin

We show here that a wide spectrum of cellular stresses activates Chk1 in a manner dependent on Claspin (**Fig. 7**). Right now, it is not clear how Claspin is involved in Chk1 activation during stress-induced responses at the molecular level. Some stresses (Ar, heat, and H_2_O_2_) may impede replication fork progression, and this may directly activate ATR-Claspin-Chk1. Others may not inhibit DNA replication, but Chk1 may be indirectly activated. By imaging and FACS-based analyses, we show that Chk1 activation in S phase depends on Claspin and that in G1 phase is largely independent of Claspin. In yeast, Mrc1, the Claspin homologue, and Rad9 are two mediator proteins that are required for checkpoint activation (phosphorylation of Rad53), though both act redundantly in Rad53 phosphorylation [49–50]. Mrc1 is required specifically for S phase replication checkpoint, while Rad9 regulates checkpoint throughout cell cycle. The roles of potential mammalian Rad9 homologue, 53BP1 or Mdc1, in Chk1 activation need to be evaluated.

**Figure 7:**
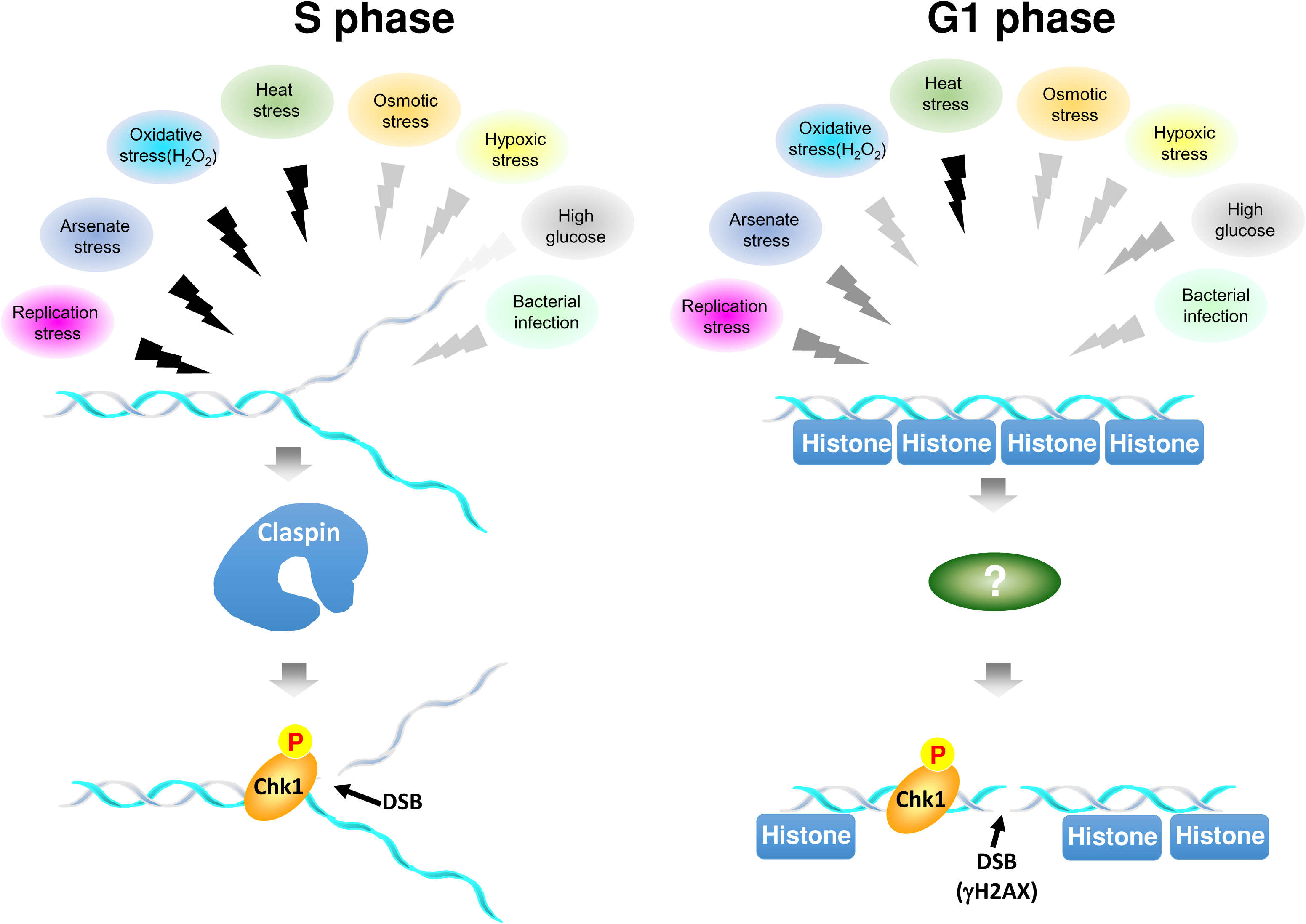
Summary on stress-mediated Chk1 activation during cell cycle. Various biological stresses activate Chk1. Overall, the Chk1 activation depends on Claspin. However, Chk1 activation is more strictly dependent on Claspin during S phase, while that in G1 phase is less dependent on Claspin. The colors of the zigzag lines represent the strength of signal, black being highest and lighter gray being lower. During S phase (left), stresses with black arrows inhibit DNA replication and may activate Chk1 and induce DNA damages at the stalled fork. Other signals also activate Chk1 albeit at a lower level. During G1 (right), all the signals can activate Chk1 to different extents. Heat, which strongly inhibits DNA replication, can vigorously activate Chk1 and DSB signal in G1 phase as well. During G1, γ−H2AX signal may represent actual DNA breaks or could be results of reorganization of chromatin structures induced by stresses.

Although single cell analyses indicate less dependency on Claspin for Chk1 activation in G1 phase, the population analyses of Chk1 activation by western analyses show that pChk1(S345) in the presence of stresses is largely dependent on Claspin (**Fig. 5A and B**). This is probably due to the fact that the level of Chk1 activation in G1 phase is generally lower than that observed in S phase (see right panel of **Figure 1E**; compare the red bar and the sum of the yellow and green bars).

Heat stress strongly inhibits DNA replication and also induces γ-H2AX signals in our experimental system. This finding leads to prediction that pChk1 and γ-H2AX signals predominantly appear during S phase. Indeed, HU, Ar or H_2_O_2_, which inhibit DNA replication, induce these signals predominantly in S phase cells. In contrast, heat treatment induces them in more than 90% G1 cells in largely Claspin-independent manner. Similarly, HG activates Chk1 and γ-H2AX in G1 phase more efficiently than in S phase cells. The activation of γ-H2AX-pChk1 in G1 phase may reflect alteration of chromatin organization or epigenome state induced by the stresses, rather than DNA damages. Alternatively, aberrant transcription induced by stresses may generate RNA-DNA hybrids that may lead to DNA damages.

A previous study showed that Mrc1, the Claspin homolog in yeast, is phosphorylated through different SAPKs upstream of Mrc1, each of which responds specifically to different stresses including osmotic, heat, oxidative stress and low glucose [38]. In mammalian cells as well, different stresses can activate Claspin via different SAPKs upstream of Claspin [51–52]. Indeed, a recent report showed that osmotic stress induced Claspin phosphorylation by activated SAPK p38 and facilitated the repair of lesions in human cells [20]. We found that Claspin undergoes hyperphosphorylation in response to various stresses, suggesting different stresses may induce differential phosphorylation of Claspin, as indicated by the distinct shifted bands.

Our findings indicate that various biological stresses activate Chk1 in both Claspin-dependent and -independent manners. They may directly interfere with DNA replication machinery or integrity of template DNA, or affects the transcription profiles as well as chromatin state, ultimately generating the sources for genomic instability. Activation of the effector kinase Chk1 may serve for protection of the genome from stress-induced lesions by modulating replication and cell cycle progression. Further studies on crosstalks between cellular responses to various biological stresses and replication checkpoint pathway would reveal novel molecular mechanisms on how cells maintain genome integrity in the face of various environmental stresses.

## Materials and Methods

### Cell lines

HeLa, U2OS, HCT116, and 293T cells were obtained from ATCC. *Claspin flox /-* Mouse Embryonic Fibroblasts (MEFs) were established from E12.5 embryos [35]. *Claspin flox /-* MEFs stably expressing the wild-type or DE/A mutant Claspin were established by infecting recombinant retroviruses expressing these cDNAs [35]. Cells were grown in Dulbecco’s modified Eagle’s medium (high glucose) supplemented with 15% fetal bovine serum (NICHIREI), 2 mM L-glutamine, 1% sodium pyruvate, 100 U/ml penicillin and 100 μg/ml streptomycin in a humidified atmosphere of 5% CO_2_, 95% air at 37°C.

### Antibodies

Antibodies used in this study are as follows. Anti-human Claspin was generated against the human recombinant Claspin with aa896–1,014 produced in *E. coli.* Anti-Chk1 phospho-S345 (#2348), anti-Chk1 phospho-S317 (#2344), anti-p44/42 MAPK (Erk1/2) (#4695), anti-SAPK/JNK (#9252), p38 MAPK (#8690), anti-p38 MAPK T180/Y182 (#4511), anti-p44/42 MAPK (Erk1/2) T202/Y204 (#4370), anti-SAPK/JNK T183/Y185 (#4668), Caspase-9 (#9508), Cleaved Caspase-3(#9661), and Mcl-1 (#5453) were obtained from Cell Signaling. Anti-α Tubulin (sc23948), anti-MCM2 (sc-9839), and anti-Chk1 (sc-8408), were obtained from Santa Cruz. Anti-phospho-H2A.X S139 (06-536) was purchased from Merck. Anti-BrdU (Ab6326) was purchased from Abcam. Anti-ATR 14hosphor-T1989 (GTX128145) was purchased from GeneTex. Anti-BrdU (555627) was purchased from BD Pharmingen. Anti-H2A.X phospho-S139 (613402), anti-Rat IgG Alexa Fluor 555 (405420), and FITC-anti-BrdU (364104) were purchased from Biolegend. RPA32 phospho-S4/S8 (A300-245A) and anti-MCM2 S53(A300-756A) was purchased from Bethyl. Anti-Mouse IgG Alexa Fluor 488 (A-11017) was purchased from Invitrogen. Goat Anti-Rabbit IgG HRP (111-035-003) and Goat Anti-Mouse IgG (115-035-003) were purchased from Jackson ImmunoResearch Laboratory.

### Claspin knockdown by siRNA

Transfection of siRNA was performed using Oligofectamine™ Transfection Reagent (Invitrogen) following manufacturer’s guidelines. All siRNAs were used at 20 pmol/ml. Transfections were performed for 48 h and cells were subjected to indicated experiments.

siRNA sequences for Claspin siRNA were as follows [53]. siClaspin-nc#7 sense GCCAAUGAUCCUUCCUUCU-TT; siClaspin-nc#7 antisense AGAAGGAAGGAUCAUUGGC-TT

### Stress conditions

To examine the stress responses in cancer cells, cells were treated with 2 mM hydroxyurea (HU), 50 J/m^2^ of UV, 100 μM Thymol, 42°C (heat shock), 50 mM NaCl, 50 μM H_2_O_2_, 2 μg/ml *E. coli* lipopolysaccharides, 400 μM Arsenate salt (Ar), 4°C (cold shock), DMEM with 30 mM glucose (high glucose), DMEM with 5.55 mM glucose (low glucose) or hypoxia [0.5% oxygen concentration in a CO_2_ incubator MG-70M (TAITEC)], respectively, for 3 hr, unless otherwise stated.

### Immunoblotting

To obtain whole cell extract (WCE), cells were first seeded in 12-well plates and cultured overnight. Exponentially growing cells were then treated with indicated biological stresses for 3 hr at 37°C. Cells were washed by PBS twice and directly resuspended by 1x sample buffer (Cold Spring Harbor Protocols). WCE was then run on 5–20% gradient SDS–polyacrylamide gel electrophoresis (PAGE; ATTO) and then transferred to Hybond ECL membranes (GE Healthcare) followed by incubation with indicated antibodies. Detection was conducted with Chemi-Lumi One Series for HRP (Nacalai) and images were obtained with LAS4000 (Fujifilm).

### Flow cytometry and cell cycle analysis

Cells were treated with indicated stresses and incubated with Bromodeoxyuridine (BrdU) at the final concentration of 20 μM for the last 15 min before the harvest. Cells were then washed and harvested. Cells were fixed with 4% PFA and incubated at 4°C overnight. Cells then were then washed by PBS supplemented with 5% BSA and permeabilized and denatured by Triton X-100 (Final concentration: 0.25%) and HCl (Final concentration: 2N), respectively. Cells were then washed and all residual acid was neutralized by 0.1M sodium borate for 2-min incubation. After wash, cells were then stained with anti-BrdU antibody conjugated with FITC and other primary antibodies diluted in wash buffer. Cells were stained with secondary antibodies at RT for 1 h. After washes, cells were then incubated with propidium iodide (PI) at RT for 30 min and samples were resuspended with PBS on ice and analyzed by flow cytometer BD LSRFortessa™ X-20. Data were then processed by FlowJo software.

### Immunostaining

FUCCI cells were treated with indicated stresses conditions for 3 hr and washed by PBS for three times. Cells were fixed with 4 %PFA in PBS for 15 min and then washed with PBS for three times. After wash, cells were permeabilized by 0.5% Triton® X-100 in PBS at RT for 20 min. After permeabilization, cells were blocked in 3 % BSA/PBS for 1 hr and indicated antibody staining. After staining, observation was performed and analyzed by Zeiss LSM780.

### DNA fiber assay

Exponentially growing cells were pulse labeled with 25 µM 5-Iodo-2’-deoxyuridine (IdU) at 37°C for 20 min. Cells were then quickly washed with PBS for three times and labeled by CldU (5-Chloro-2’-deoxyuridine) at 37°C for 20 min with indicated biological stresses. Cells were then incubated with 2.5mM thymidine at RT for 30sec after quick washes with PBS three times. Cells were then trypsinized and resuspended with PBS at the cell density of 1x10^6^ cells/ml. 2 µl of labeled cells were mixed with unlabeled cells at the ratio of 1:1 and dropped onto the slides (Pro-01; Matsunami). The cell mixture was then lysed with the buffer (200 mM Tris-HCl–50 mM EDTA with 0.5% SDS) for 5 min. Slides were tilted on the lid of a multi-well plate and DNA fibers flowed down along the slides at a constant speed. Fibers were then fixed with the solution containing methanol and acetic acid at the mix ratio of 3:1 at 4°C overnight. Fibers were then denatured by 2.5 N HCl and blocked with PBS supplemented with 3 % BSA and 0.1 % Tween20. Samples were then stained with anti-BrdU antibody [Clone: BU1/75 (ICR1); Abcam] and anti-BrdU antibody (Clone: 3D4; BD) at RT for 1 h in the dark. After incubation with primary antibodies, fibers were then incubated with high salt buffer (28mM Tris-HCl pH8.0, 500mM NaCl. 0.5% Triton X-100) at RT for 10min in the dark. Fibers were then subjected to secondary antibody reactions and Hoechst staining at RT for 1 h in the dark. Fibers were visualized with Keyence BZ-X700 and quantified and calculated by ImageJ.

## Acknowledgments

We thank the members of our laboratory for useful discussion.

## Author Contributions

H.M., C-C.Y. and H-W.H. conceived the research plans. H-W.H. and C-C.Y. conducted the experiments. H-W.H., C-CY and H.M. wrote the paper.

---

**Supplementary Table 1.**

|  | Signals | Control | HU<br>(replication stress) | Ar (arsenate) | NaCl<br>(osmotic stress) | LPS<br>(bacterial infection) | Hypoxia<br>(hypoxic stress) | Heat (high temperature stress) | H2O2<br>(oxidative stress) | HG (high glucose stress) |
| --- | --- | --- | --- | --- | --- | --- | --- | --- | --- | --- |
| Imaging | gH2AX (DNA damage) | S=G1 | S>G1 | S>G1 | S>G1 | S=G1 | S>G1 | S<G1 | S>G1 | S<G1 |
|  | pChk1 (S345)<br>(replication stress) | S=G1 | S>G1 | S=G1 | S=G1 | S=G1 | S>G1 | S<G1 | S>G1 | S<G1 |
| FACS | gH2AX (DNA damage) | - | +++ | ++ | + | + | + | +++ | +++ | + |
|  | pRPA (DNA damage) | - | +++ | ++ | + | + | + | ++ | +++ | + |
|  | pChk1 (S345)<br>(replication stress) | - | +++ | ++ | + | + | + | +++ | +++ | + |
|  | BrdU (DNA replication) | +++ | - | - | ++ | ++ | +++ | - | - | ++ |

## References

1. Gaillard, H., García-Muse, T., & Aguilera, A. (2015). Replication stress and cancer. Nature Reviews Cancer, 15(5), 276–289. https://doi.org/10.1038/nrc3916

2. Zeman, M. K., & Cimprich, K. A. (2014). Causes and consequences of replication stress. Nature cell biology, 16(1), 2–9. https://doi.org/10.1038/ncb2897

3. Yang, C. C., Kato, H., Shindo, M., & Masai, H. (2019). Cdc7 activates replication checkpoint by phosphorylating the Chk1-binding domain of Claspin in human cells. Elife, 8, e50796. https://doi.org/10.7554/eLife.50796

4. Hsiao, H. W., Yang, C. C., & Masai, H. (2021). Roles of Claspin in regulation of DNA replication, replication stress responses and oncogenesis in human cells. Genome Instability & Disease, 2(5), 263–280. https://doi.org/10.1007/s42764-021-00049-8

5. Chini, C. C. S., Wood, J., & Chen, J. (2006). Chk1 is required to maintain claspin stability. Oncogene, 25(30), 4165–4171. https://doi.org/10.1038/sj.onc.1209447

6. Kumagai, A., & Dunphy, W. G. (2000). Claspin, a novel protein required for the activation of Chk1 during a DNA replication checkpoint response in Xenopus egg extracts. Molecular cell, 6(4), 839–849. https://doi.org/10.1016/S1097-2765(05)00092-4

7. Lee, J., Kumagai, A., & Dunphy, W. G. (2003). Claspin, a Chk1-regulatory protein, monitors DNA replication on chromatin independently of RPA, ATR, and Rad17. Molecular cell, 11(2), 329–340. https://doi.org/10.1016/S1097-2765(03)00045-5

8. Chini CC., & Chen J. (2006) Repeated phosphopeptide motifs in human claspin are phosphorylated by Chk1 and mediate claspin function. Journal of Biological Chemistry 281:33276–33282. https://doi.org/10.1074/jbc.M604373200

9. Yoo HY., Jeong SY., & Dunphy WG. (2006) Site-specific phosphorylation of a checkpoint mediator protein controls its responses to different DNA structures. Genes & Development 20:772–783. https://doi.org/10.1101/gad.1398806

10. Lindsey-Boltz LA., Serçin Özdemirhan., Choi J-H., & Sancar A. (2009) Reconstitution of Human Claspin-mediated Phosphorylation of Chk1 by the ATR (Ataxia Telangiectasia-mutated and Rad3-related) Checkpoint Kinase. Journal of Biological Chemistry 284:33107–33114. https://doi.org/10.1074/jbc.M109.064485

11. Alcasabas AA., Osborn AJ., Bachant J., Hu F., Werler PJ., Bousset K., Furuya K., Diffley JF., Carr AM., & Elledge SJ.(2001) Mrc1 transduces signals of DNA replication stress to activate Rad53. Nature Cell Biology 3:958–965. https://doi.org/10.1038/ncb1101-958

12. Osborn AJ., & Elledge SJ. (2003) Mrc1 is a replication fork component whose phosphorylation in response to DNA replication stress activates Rad53. Genes & Development 17:1755–1767. https://doi.org/10.1101/gad.1098303

13. Tanaka K., & Russell P. (2001) Mrc1 channels the DNA replication arrest signal to checkpoint kinase Cds1. Nature Cell Biology 3:966–972. https://doi.org/10.1038/ncb1101-966

14. Meng Z., Capalbo L.,Glover DM., & Dunphy WG.(2011) Role for casein kinase 1 in the phosphorylation of claspin on critical residues necessary for the activation of Chk1 Molecular Biology of the Cell 22:2834–2847. https://doi.org/10.1091/mbc.e11-01-0048

15. Elvers, I., Hagenkort, A., Johansson, F., Djureinovic, T., Lagerqvist, A., Schultz, N., … & Helleday, T. (2012). CHK1 activity is required for continuous replication fork elongation but not stabilization of post-replicative gaps after UV irradiation. Nucleic acids research, 40(17), 8440–8448. https://doi.org/10.1093/nar/gks646

16. Furusawa, Y., Iizumi, T., Fujiwara, Y., Zhao, Q. L., Tabuchi, Y., Nomura, T., & Kondo, T. (2012). Inhibition of checkpoint kinase 1 abrogates G2/M checkpoint activation and promotes apoptosis under heat stress. Apoptosis, 17(1), 102–112. https://doi.org/10.1007/s10495-011-0660-7

17. Ramachandran, S., Ma, T. S., Griffin, J., Ng, N., Foskolou, I. P., Hwang, M. S., … & Hammond, E. M. (2021). Hypoxia-induced SETX links replication stress with the unfolded protein response. Nature Communications, 12(1), 1–14. https://doi.org/10.1038/s41467-021-24066-z

18. Yang, Y., Liu, C., Xie, T., Wang, D., Chen, X., Ma, L., & Zhang, A. (2021). Role of inhibiting Chk1-p53 pathway in hepatotoxicity caused by chronic arsenic exposure from coal-burning. Human & Experimental Toxicology, 40(7), 1141–1152. https://doi.org/10.1177/0960327120988880

19. Dmitrieva, N. I., Cai, Q., & Burg, M. B. (2004). Cells adapted to high NaCl have many DNA breaks and impaired DNA repair both in cell culture and in vivo. Proceedings of the National Academy of Sciences, 101(8), 2317–2322. https://doi.org/10.1073/pnas.0308463100

20. Ulsamer, A., Martínez-Limón, A., Bader, S., Rodríguez-Acebes, S., Freire, R., Méndez, J., … & Posas, F. (2022). Regulation of Claspin by the p38 stress-activated protein kinase protects cells from DNA damage. Cell Reports, 40(12), 111375., https://doi.org/10.1016/j.celrep.2022.111375.

21. Sharma, K., Kumar, C., Kéri, G., Breitkopf, S. B., Oppermann, F. S., & Daub, H. (2010). Quantitative analysis of kinase-proximal signaling in lipopolysaccharide-induced innate immune response. Journal of proteome research, 9(5), 2539–2549. https://doi.org/10.1021/pr901192p

22. Zhong, A., Chang, M., Yu, T., Gau, R., Riley, D. J., Chen, Y., & Chen, P. L. (2018). Aberrant DNA damage response and DNA repair pathway in high glucose conditions. Journal of cancer research updates, 7(3), 64. https://doi.org/10.6000/1929-2279.2018.07.03.1

23. Kumar, V., Agrawal, R., Pandey, A., Kopf, S., Hoeffgen, M., Kaymak, S., … & Nawroth, P. P. (2020). Compromised DNA repair is responsible for diabetes-associated fibrosis. The EMBO journal, 39(11), e103477.

24. Willis, J., Patel, Y., Lentz, B. L., & Yan, S. (2013). APE2 is required for ATR-Chk1 checkpoint activation in response to oxidative stress. Proceedings of the National Academy of Sciences, 110(26), 10592–10597. https://doi.org/10.1073/pnas.1301445110

25. Tam, L. M., Price, N. E., & Wang, Y. (2020). Molecular mechanisms of arsenic-induced disruption of DNA repair. Chemical research in toxicology, 33(3), 709–726. https://doi.org/10.1021/acs.chemrestox.9b00464

26. Joe, Y., Jeong, J. H., Yang, S., Kang, H., Motoyama, N., Pandolfi, P. P., … & Kim, M. K. (2006). ATR, PML, and CHK2 play a role in arsenic trioxide-induced apoptosis. Journal of Biological Chemistry, 281(39), 28764–28771. https://doi.org/10.1074/jbc.M604392200

27. Taniuchi, S., Miyake, M., Tsugawa, K., Oyadomari, M., & Oyadomari, S. (2016). Integrated stress response of vertebrates is regulated by four eIF2α kinases. Scientific reports, 6(1), 1–11. https://doi.org/10.1038/srep32886

28. Martin, L., Rainey, M., Santocanale, C., & Gardner, L. B. (2012). Hypoxic activation of ATR and the suppression of the initiation of DNA replication through cdc6 degradation. Oncogene, 31(36), 4076–4084. https://doi.org/10.1038/onc.2011.585

29. Pires, I. M., Bencokova, Z., McGurk, C., & Hammond, E. M. (2010). Exposure to acute hypoxia induces a transient DNA damage response which includes Chk1 and TLK1. Cell cycle, 9(13), 2502–2507. https://doi.org/10.4161/cc.9.13.12059

30. Pires, I. M., Bencokova, Z., Milani, M., Folkes, L. K., Li, J. L., Stratford, M. R., … & Hammond, E. M. (2010). Effects of Acute versus Chronic Hypoxia on DNA Damage Responses and Genomic Instability. Cancer research, 70(3), 925–935. https://doi.org/10.1158/0008-5472.CAN-09-2715

31. Foskolou, I. P., Jorgensen, C., Leszczynska, K. B., Olcina, M. M., Tarhonskaya, H., Haisma, B., … & Hammond, E. M. (2017). Ribonucleotide reductase requires subunit switching in hypoxia to maintain DNA replication. Molecular cell, 66(2), 206–220. https://doi.org/10.1016/j.molcel.2017.03.005

32. Ng, N., Purshouse, K., Foskolou, I. P., Olcina, M. M., & Hammond, E. M. (2018). Challenges to DNA replication in hypoxic conditions. The FEBS journal, 285(9), 1563–1571. https://doi.org/10.1111/febs.14377

33. Bolland, H., Ma, T. S., Ramlee, S., Ramadan, K., & Hammond, E. M. (2021). Links between the unfolded protein response and the DNA damage response in hypoxia: A systematic review. Biochemical Society Transactions, 49(3), 1251–1263. https://doi.org/10.1042/BST20200861

34. Cabrera, E., Hernández-Pérez, S., Koundrioukoff, S., Debatisse, M., Kim, D., Smolka, M. B., … & Gillespie, D. A. (2017). PERK inhibits DNA replication during the Unfolded Protein Response via Claspin and Chk1. Oncogene, 36(5), 678–686. https://doi.org/10.1038/onc.2016.239

35. Bader, S. B., Ma, T. S., Simpson, C. J., Liang, J., Maezono, S. E. B., Olcina, M. M., … & Hammond, E. M. (2021). Replication catastrophe induced by cyclic hypoxia leads to increased APOBEC3B activity. Nucleic acids research, 49(13), 7492–7506. https://doi.org/10.1093/nar/gkab551

36. Casalino-Matsuda, S. M., Berdnikovs, S., Wang, N., Nair, A., Gates, K. L., Beitel, G. J., & Sporn, P. H. (2021). Hypercapnia selectively modulates LPS-induced changes in innate immune and DNA replication-related gene transcription in the macrophage. Interface focus, 11(2), 20200039.

37. Chen, M. Y., Hsu, W. C., Hsu, S. C., Yang, Y. S., Chuang, T. H., Lin, W. J., … & Su, Y. W. (2019). PP4 deficiency leads to DNA replication stress that impairs immunoglobulin class switch efficiency. Cell Death & Differentiation, 26(7), 1221–1234. https://doi.org/10.1038/s41418-018-0199-z

38. Tuul, M., Kitao, H., Iimori, M., Matsuoka, K., Kiyonari, S., Saeki, H., … & Maehara, Y. (2013). Rad9, Rad17, TopBP1 and claspin play essential roles in heat-induced activation of ATR kinase and heat tolerance. PLoS One, 8(2), e55361. https://doi.org/10.1371/journal.pone.0055361

39. Luo, Z., Zheng, K., Fan, Q., Jiang, X., & Xiong, D. (2017). Hyperthermia exposure induces apoptosis and inhibits proliferation in HCT116 cells by upregulating miR-34a and causing transcriptional activation of p53. Experimental and Therapeutic Medicine, 14(6), 5379–5386. https://doi.org/10.3892/etm.2017.5257

40. Duch, A., Canal, B., Barroso, S. I., García-Rubio, M., Seisenbacher, G., Aguilera, A., … & Posas, F. (2018). Multiple signaling kinases target Mrc1 to prevent genomic instability triggered by transcription-replication conflicts. Nature communications, 9(1), 1–14. https://doi.org/10.1038/s41467-017-02756-x

41. Coluzzi, E., Leone, S., & Sgura, A. (2019). Oxidative stress induces telomere dysfunction and senescence by replication fork arrest. Cells, 8(1), 19. https://doi.org/10.3390/cells8010019

42. Somyajit, K., Gupta, R., Sedlackova, H., Neelsen, K. J., Ochs, F., Rask, M. B., … & Lukas, J. (2017). Redox-sensitive alteration of replisome architecture safeguards genome integrity. Science, 358(6364), 797–802. https://doi.org/10.1126/science.aao3172

43. Ciminera, A. K., Shuck, S. C., & Termini, J. (2021). Elevated glucose increases genomic instability by inhibiting nucleotide excision repair. Life science alliance, 4(10). https://doi.org/10.26508/lsa.202101159

44. Hu, C. M., Tien, S. C., Hsieh, P. K., Jeng, Y. M., Chang, M. C., Chang, Y. T., Chen, Y. J., Chen, Y. J., Lee, E. Y. P., & Lee, W. H. (2019). High Glucose Triggers Nucleotide Imbalance through O-GlcNAcylation of Key Enzymes and Induces KRAS Mutation in Pancreatic Cells. Cell metabolism, 29(6), 1334–1349.e10. https://doi.org/10.1016/j.cmet.2019.02.005

45. Yang, C. C., & Masai, H. (2022). Claspin is required for growth recovery from serum starvation through regulating the PI3K-PDK1-mTOR pathway. bioRxiv. https://doi.org/10.1101/2022.01.21.475743

46. Yang, C. C., Suzuki, M., Yamakawa, S., Uno, S., Ishii, A., Yamazaki, S., … & Masai, H. (2016). Claspin recruits Cdc7 kinase for initiation of DNA replication in human cells. Nature communications, 7(1), 1–14. https://doi.org/10.1038/ncomms12135

47. Sakaue-Sawano, A., Kurokawa, H., Morimura, T., Hanyu, A., Hama, H., Osawa, H., … & Miyawaki, A. (2008). Visualizing spatiotemporal dynamics of multicellular cell-cycle progression. Cell, 132(3), 487–498.

48. Im, J. S., & Lee, J. K. (2008). ATR-dependent activation of p38 MAP kinase is responsible for apoptotic cell death in cells depleted of Cdc7. Journal of Biological Chemistry, 283(37), 25171–25177. https://doi.org/10.1074/jbc.M802851200

49. Bacal, J., Moriel-Carretero, M., Pardo, B., Barthe, A., Sharma, S., Chabes, A., … & Pasero, P. (2018). Mrc1 and Rad9 cooperate to regulate initiation and elongation of DNA replication in response to DNA damage. The EMBO journal, 37(21), e99319. https://doi.org/10.15252/embj.201899319

50. McClure, A. W., & Diffley, J. F. (2021). Rad53 checkpoint kinase regulation of DNA replication fork rate via Mrc1 phosphorylation. Elife, 10, e69726. https://doi.org/10.7554/eLife.69726

51. Llopis, A., Salvador, N., Ercilla, A., Guaita-Esteruelas, S., Barrantes, I. D. B., Gupta, J., … & Agell, N. (2012). The stress-activated protein kinases p38α/β and JNK1/2 cooperate with Chk1 to inhibit mitotic entry upon DNA replication arrest. Cell Cycle, 11(19), 3627–3637. https://doi.org/10.4161/cc.21917

52. Canovas, B., & Nebreda, A. R. (2021). Diversity and versatility of p38 kinase signalling in health and disease. Nature Reviews Molecular Cell Biology, 22(5), 346–366. https://doi.org/10.1038/s41580-020-00322-w

53. Uno, S., & Masai, H. (2011). Efficient expression and purification of human replication fork-stabilizing factor, Claspin, from mammalian cells: DNA-binding activity and novel protein interactions. Genes to Cells, 16(8), 842–856. https://doi.org/10.1111/j.1365-2443.2011.01535.

